# Metabolic transitions reprogram translation through sequence-dependent regulation

**DOI:** 10.64898/2026.11.15.766763

**Authors:** Xavier Hernandez-Alias, Selay Kaya, Konstantin Doychev, Felix Schroeder, Bohdana Hermann, Berra Madran, Sina Majidian, Julien Gagneur, Danny D. Nedialkova

## Abstract

Cells suppress protein synthesis during starvation, but how translation recovers upon nutrient repletion remains poorly understood. To address this gap, we followed translational recovery in budding yeast after repletion of glucose, nitrogen, or phosphate, combining time-resolved metabolomics, structural proteomics, transcriptomics, and spike-in-calibrated ribosome profiling. Nutrient addition rapidly altered ribosome occupancy in distinct, transcript-specific programs. To identify their sequence determinants, we developed RiboNet, a deep learning framework that predicts ribosome load and positions along transcripts from mRNA sequence. RiboNet-guided *in silico* mutagenesis revealed reduced pausing at serine codons upon glucose repletion and transiently increased pausing at leucine codons following nitrogen repletion. These responses coincided with altered cognate tRNA charging and metabolite-associated structural changes in aminoacyl-tRNA synthetases. RiboNet also identified a conserved 5′ UTR motif, GAUY, that increased reporter protein output following glucose repletion. Our findings reveal how metabolic state alters the contributions of mRNA sequence features to translation initiation and elongation.

## INTRODUCTION

Nutrient availability is a major determinant of cellular growth, requiring continuous coordination between metabolism and gene expression. In the budding yeast *Saccharomyces cerevisiae*, carbon, nitrogen, and phosphate are sensed through largely distinct networks, acting through protein kinase A, TORC1, and the PHO pathway, respectively^1–6^. Protein synthesis is particularly sensitive to nutrient limitation because of its substantial energetic demands^7^. Glucose deprivation causes a rapid and widespread shutdown of protein synthesis that is reversed upon glucose repletion^8,9^, involving reduced translation initiation, slower elongation, and changes in mRNA fate^10–13^. The fastest known component of this response is metabolic: ATP falls within seconds of glucose removal, below the affinity of initiation factors such as eIF4A, so that initiation stops on pre-existing mRNAs^14^. Because this mechanism senses energy rather than glucose itself, it is unclear whether other nutrient stresses act through shared or distinct translational controls. Nitrogen starvation similarly represses translation through TORC1 and alters the abundance of translation-related factors^15–17^.

Although translational suppression during starvation has been extensively studied, far less is known about how translation is restored when a nutrient returns, and previous work on repletion has focused largely on metabolic and transcriptional responses^18–20^. Recovery is not simply the reversal of shutdown. Most mRNAs released from polysomes during glucose withdrawal are degraded, and fewer than 400 can be reactivated without new transcription^12^. How ribosomes are redistributed across individual mRNAs during recovery, and whether the sequence features that distinguish transcripts under steady-state conditions operate in the same way during an abrupt transition, remains unknown.

Resolving the temporal relationship between metabolic adaptation, transcription, and translation requires quantitative measurements during the early stages of nutrient repletion. Metabolite concentrations can change within seconds^8,14,21^, potentially affecting the translational machinery before substantial changes in mRNA abundance occur. Ribosome profiling provides nucleotide-resolution measurements of ribosome occupancy^22^, but conventional library normalization can obscure global changes in translation. Spike-in calibration enables estimates of absolute ribosome numbers per mRNA^23^, while distinct footprint lengths provide information about ribosomal states during elongation^24,25^. Interpreting these measurements is challenging because ribosome occupancy at individual codons depends on both ribosome load (the average number of ribosomes per mRNA) and how quickly ribosomes move along the coding sequence. Existing machine-learning approaches generally model these properties separately, predicting either transcript-level output from the 5′ UTR^26,27^ or local ribosome density along coding sequences^28–31^, and few explicitly model how these properties change across cellular state^28,30^. By treating ribosome numbers and their positional distribution separately, these approaches are limited in their ability to capture the interdependence of initiation and elongation^32–36^. Recent work suggests that elongation rates remain nearly constant across steady-state nutrient limitations, while ribosome and mRNA concentrations scale with growth rate^37,38^. Whether this stability persists during rapid changes in nutrient availability, or whether individual transcripts exhibit distinct elongation responses, remains unknown.

Here, we examined translational recovery following glucose, nitrogen, and phosphate repletion in *S. cerevisiae* through time-resolved measurements of cellular metabolites, protein structure, mRNA abundance, and ribosome occupancy. These analyses revealed substantial differences in the timing and magnitude of translational responses between nutrients and individual transcripts. To investigate their sequence determinants, we developed RiboNet, which jointly models the number of ribosomes on an mRNA and their distribution along its coding sequence across metabolic conditions. Model interpretation traced the most prominent codon-level changes to the A-site codon (serine codons after glucose repletion and leucine codons after nitrogen repletion), and these changes tracked cognate tRNA charging and coincided with structural changes in the corresponding aminoacyl-tRNA synthetases. RiboNet also uncovered GAUY, a conserved 5′ UTR motif whose effects on protein output during glucose repletion were experimentally validated. Together, our findings reveal how metabolic state influences sequence-dependent regulation of translation initiation and elongation.

## RESULTS

### Nutrient repletion triggers rapid, nutrient-specific responses

To characterize the early cellular response to nutrient repletion, we established a time-resolved experimental system in budding yeast (**Fig. 1A**, see *Methods*). We avoided confounding effects of auxotrophy on gene expression^39^ by using the prototrophic *S288c*-derived strain YSBN6. Cells were grown overnight in synthetic defined (SD) medium or in nitrogen– or phosphate-limiting medium to exhaust the large intracellular stores that would otherwise buffer a short deprivation^40^, and then deprived of glucose, nitrogen, or phosphate for 1 h (**Extended Data Fig. 1A**, see *Methods*). Compared with cells growing in SD medium, nitrogen– and phosphate-deprived cells showed extensive, largely overlapping proteomic changes, including a lower abundance of ribosome biogenesis proteins and a higher abundance of proteins involved in organonitrogen catabolism and glutamine-family amino acid metabolism (**Extended Data Fig. 1** and **Supplementary Table 1**). Because only these two conditions included overnight limitation, part of this shared response may reflect the longer pretreatment. Glucose deprivation altered fewer proteins overall and induced hexose transporters (**Extended Data Fig. 1** and **Supplementary Table 1**). All three nutrient deprivations increased the abundance of proteins annotated to organic acid metabolism (**Extended Data Fig. 1**).

**Fig. 1.**
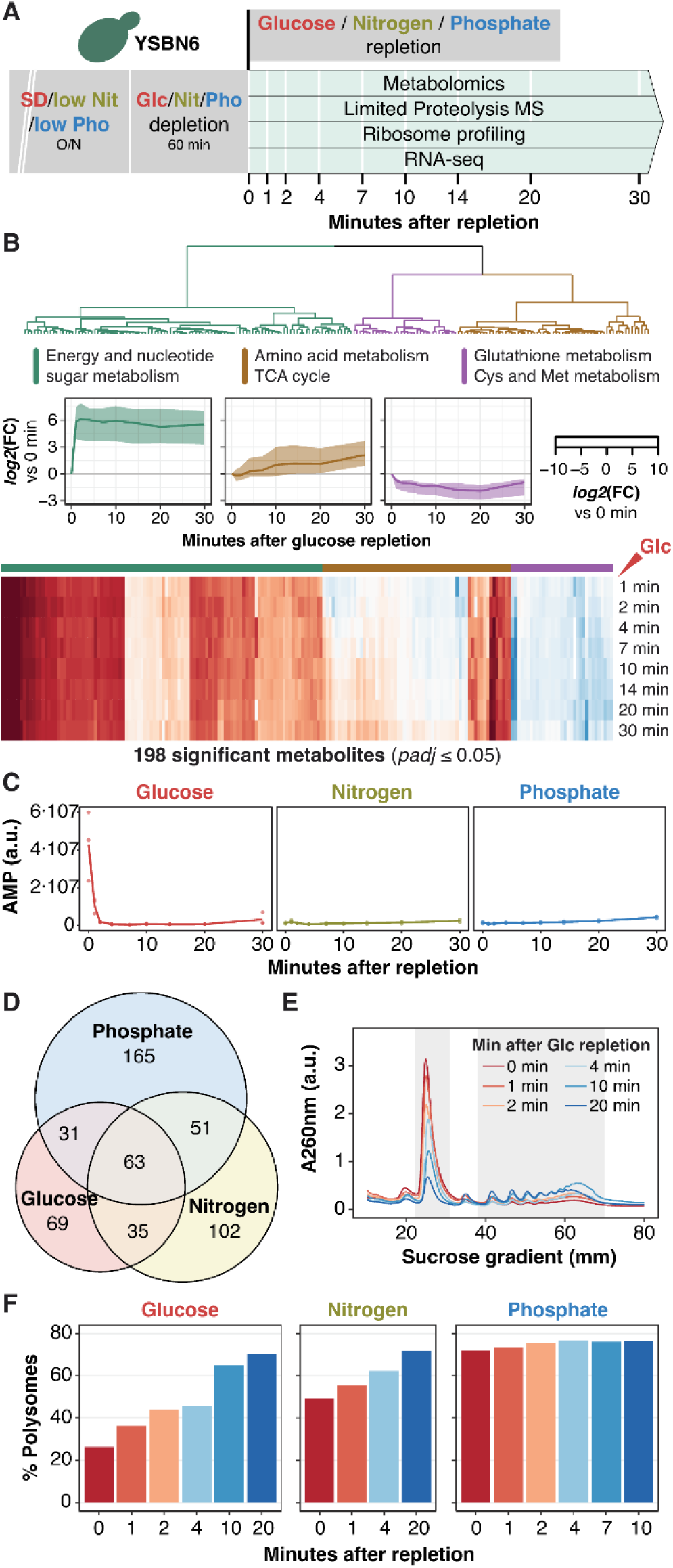
Nutrient repletion triggers rapid, nutrient-specific responses. (A) Visual representation of the multi-omic time-resolved measurements of glucose (Glc), nitrogen (Nit) and phosphate (Pho) repletions. (B) Clustering analysis of the 198 metabolic features significantly changing during glucose repletion. Significant features were identified using *limma* with a cubic spline (n=27, 3 replicates × 9 time points, *p_adj_* ≤ 0.05). Metabolites were clustered by time pattern using 1 − *R_Pearson_* as distance metric and Ward’s minimum variance linkage method. For each cluster, the line plot depicts the median and the shaded interquartile range (IQR), as well as the top hits of the corresponding peak set enrichment analysis of metabolic pathways (Fisher’s exact test, *p* ≤ 0.05). Heatmap shows the average of n=3 biological replicates. (C) AMP abundance over the 30-minute time course for each nutrient repletion. Lines show the average of n=3 biological replicates. (D) Venn diagram of significantly changing metabolite features along the 30-minute repletion for each of the three nutrients. (E) Polysome profiles upon glucose repletion (n=1 per time point). The shaded area indicates the monosomes and polysomes (≥3 ribosomes per mRNA). (F) The percentage of polysomes was quantified as the area under the curve (AUC) of their profiles divided by the total AUC. Colors correspond to curves in Fig. 1E.

We then restored each nutrient and monitored metabolite changes by untargeted LC-MS/MS over 30 min (**Fig. 1A**). Nutrient repletion elicited immediate and distinct metabolic responses. In each condition, clustering the significantly changing features by time revealed three recurring patterns: features that rose sharply within the first minute and then plateaued, features that increased gradually over 30 min, and features that declined. The immediate response was strongest after glucose repletion, where the fast cluster was enriched for features mapping to energy and nucleotide sugar metabolism, consistent with immediate glucose uptake and phosphorylation; features mapping to amino acid metabolism and the TCA cycle rose gradually, and those mapping to glutathione and cysteine/methionine metabolism declined (**Fig. 1B**). After nitrogen repletion, features mapping to aromatic amino acid biosynthesis rose rapidly, and those mapping to amino acid and glutathione metabolism rose gradually, whereas features mapping to the TCA cycle and pyruvate metabolism declined (**Extended Data Fig. 2B**). After phosphate repletion, energy and nucleotide sugar features rose rapidly, those mapping to cysteine/methionine and purine metabolism increased gradually, and those mapping to glutathione and thiamine metabolism declined (**Extended Data Fig. 2C**). Of the 516 features that changed in at least one condition, 63 changed in all three and 336 in only one (**Fig. 1D**). Thus, each repletion engages a largely nutrient-specific metabolic program.

We next examined changes in nucleotide metabolism upon nutrient repletion. AMP levels were much higher in glucose-deprived cells than in nitrogen– or phosphate-deprived cells and declined to comparable levels within minutes of glucose repletion, whereas they remained low throughout nitrogen and phosphate repletion (**Fig. 1C** and **Supplementary Table 1**). This glucose-specific response indicates a rapid reversal of the adenylate imbalance associated with energy depletion. Guanosine was elevated in both glucose– and nitrogen-deprived cells relative to phosphate-deprived cells, but declined rapidly only after nitrogen repletion (**Extended Data Fig. 2A**). These observations reveal distinct, nutrient-dependent changes in purine metabolism during recovery, with the most pronounced adenylate response occurring after glucose repletion.

Nitrogen repletion predominantly affected amino acid pools and their metabolic precursors, while phosphate repletion was characterized by changes in nucleotide-related metabolites (**Extended Data Fig. 2B-C**). The limited overlap of significantly changing metabolites across the three conditions further highlighted the distinct metabolic responses to repletion of each nutrient (**Fig. 1D**).

Next, we examined translation by polysome profiling. In glucose-deprived cells, the polysome fraction was only ∼25%, consistent with the rapid inhibition of translation initiation upon glucose withdrawal^8^. The polysome fraction increased within 1 min of glucose repletion and reached ∼65–70% by 10-20 min (**Fig. 1E, F**). Nitrogen-deprived cells had an intermediate polysome fraction of ∼50% that rose to ∼70% within 20 min (**Fig. 1F; Extended Data Fig. 2D**), whereas phosphate-deprived cells maintained ∼70–75% polysomes with little change after repletion (**Fig. 1F; Extended Data Fig. 2D**). Bulk translation is therefore strongly repressed by glucose deprivation and more mildly by nitrogen deprivation, and recovers over 10-20 min after repletion.

**Extended Data Fig. 1.**
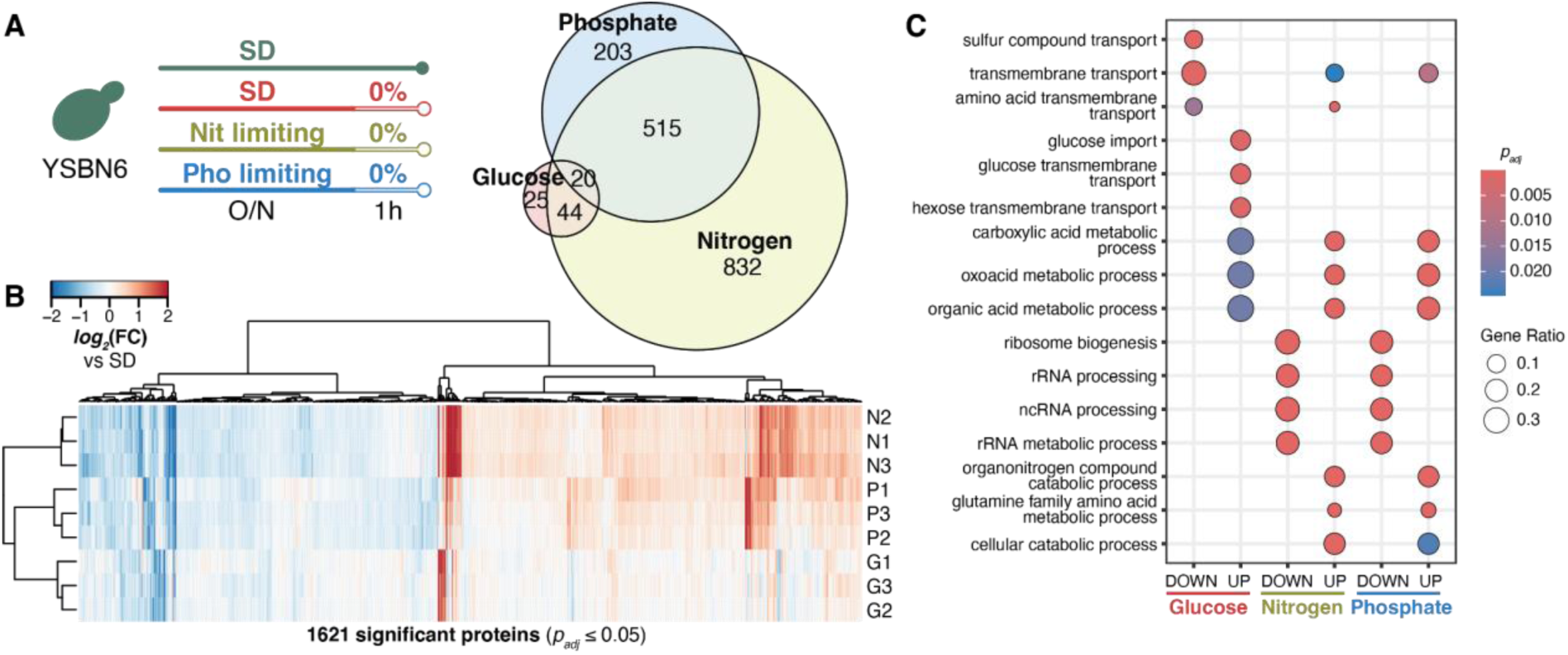
Proteomic characterization of nutrient-depleted yeast cultures. (A) Total proteomics of the starting deprived state right before nutrient repletion. Venn diagram shows significantly different proteins compared to control cells grown in synthetically defined (SD) medium. Significant proteins identified using *limma* (n=12, 3 replicates × 4 conditions, *p_adj_* ≤ 0.05). The more extensive proteomic changes observed under nitrogen and phosphate deprivation may partly reflect their longer pretreatment protocols. (B) Clustering analysis of all differentially abundant proteins in either of the three nutrient deprivations compared to yeast growing in SD medium. Hierarchical clustering uses the Euclidean distance metric and Ward’s minimum variance linkage method. (C) GO enrichment analysis of up– and down-regulated proteins in each of the three nutrient deprivations, showing the top 3 most significant biological processes (Fisher’s exact test, *p_adj_* ≤ 0.05).

**Extended Data Fig. 2.**
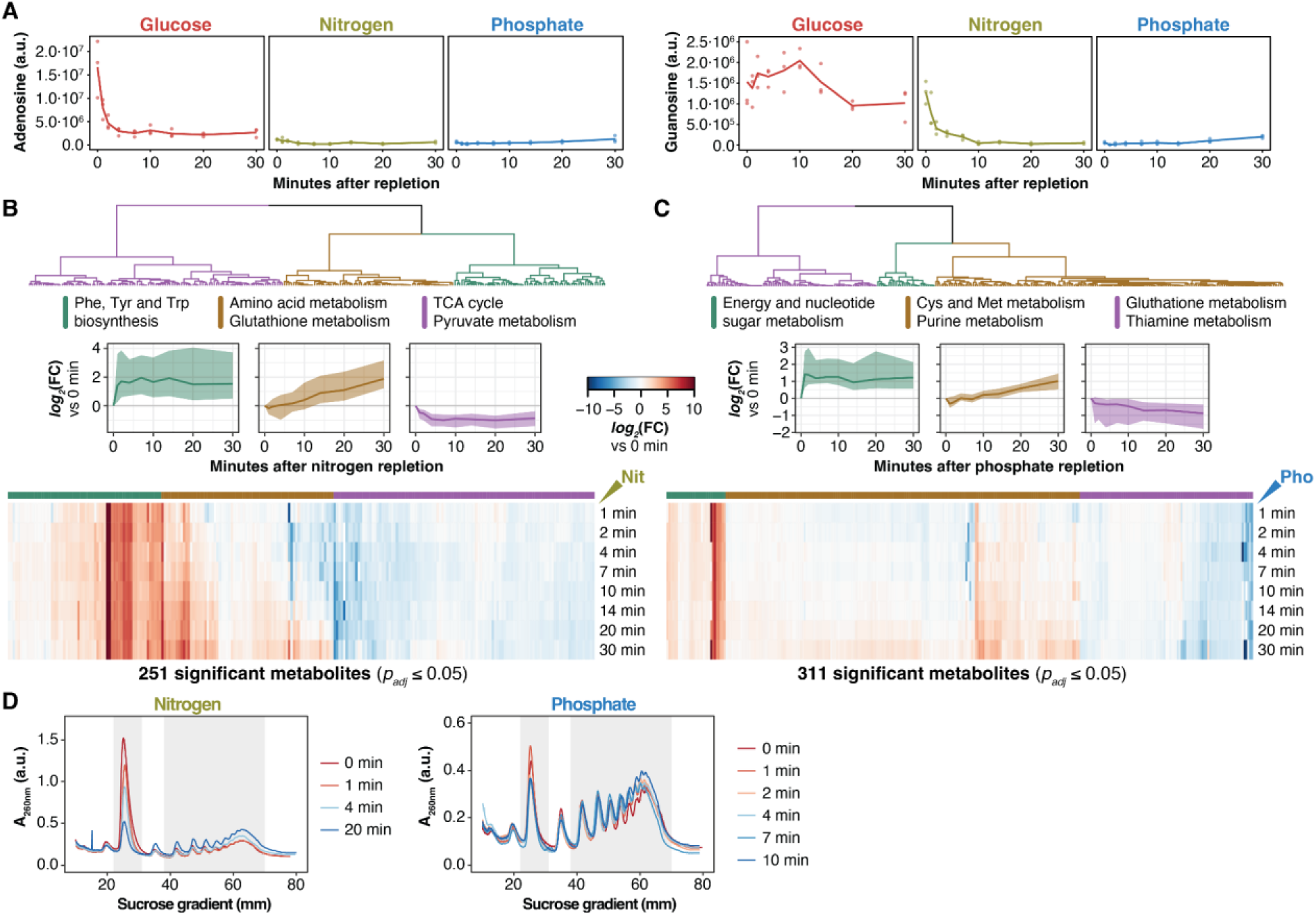
Metabolic reprogramming upon nutrient repletion induces fast translation reactivation. (A) Adenosine (left) and guanosine (right) abundance along the 30-minutes time-course for each nutrient repletion. Lines show the average of n=3 biological replicates. (B-C) Clustering analysis of all metabolic features significantly changing along the nitrogen and phosphate repletions. Significant features identified using *limma* with a cubic spline (n=27, *p_adj_* ≤ 0.05). Metabolites were clustered by time pattern using 1 − *R_Pearson_* as the distance metric and Ward’s minimum variance linkage method. For each cluster, the line plot depicts the median and the shaded IQR, as well as the top hits of the corresponding peak set enrichment analysis of metabolic pathways (Fisher’s exact test, *p* ≤ 0.05). Heatmap shows the average of n=3 biological replicates. (D) Representative polysome profiles of nitrogen and phosphate repletions (n=1 per time point). The shaded area indicates the monosomes and polysomes (≥3 ribosomes per mRNA).

### Nutrient repletion alters mRNA-specific ribosome occupancy and its relationship to codon decoding

We next asked whether early translational changes simply follow mRNA abundance. To this end, we performed ribosome profiling and matched RNA-seq at six time points over the first 10 min after nutrient repletion. Glucose repletion changed the abundance of ribosome-protected fragments (RPF) for 3,913 transcripts, of which 2,763 did not change significantly in mRNA abundance (**Fig. 2A** and **Supplementary Table 2**). After nitrogen repletion, most changes were shared between RPFs and mRNA abundance (3,286 transcripts), and phosphate repletion altered comparatively fewer transcripts in either measurement (**Fig. 2A** and **Supplementary Table 2**). These data suggest that the early glucose response acts largely at the translational level, whereas the nitrogen response is largely accompanied by changes in mRNA abundance.

**Fig. 2.**
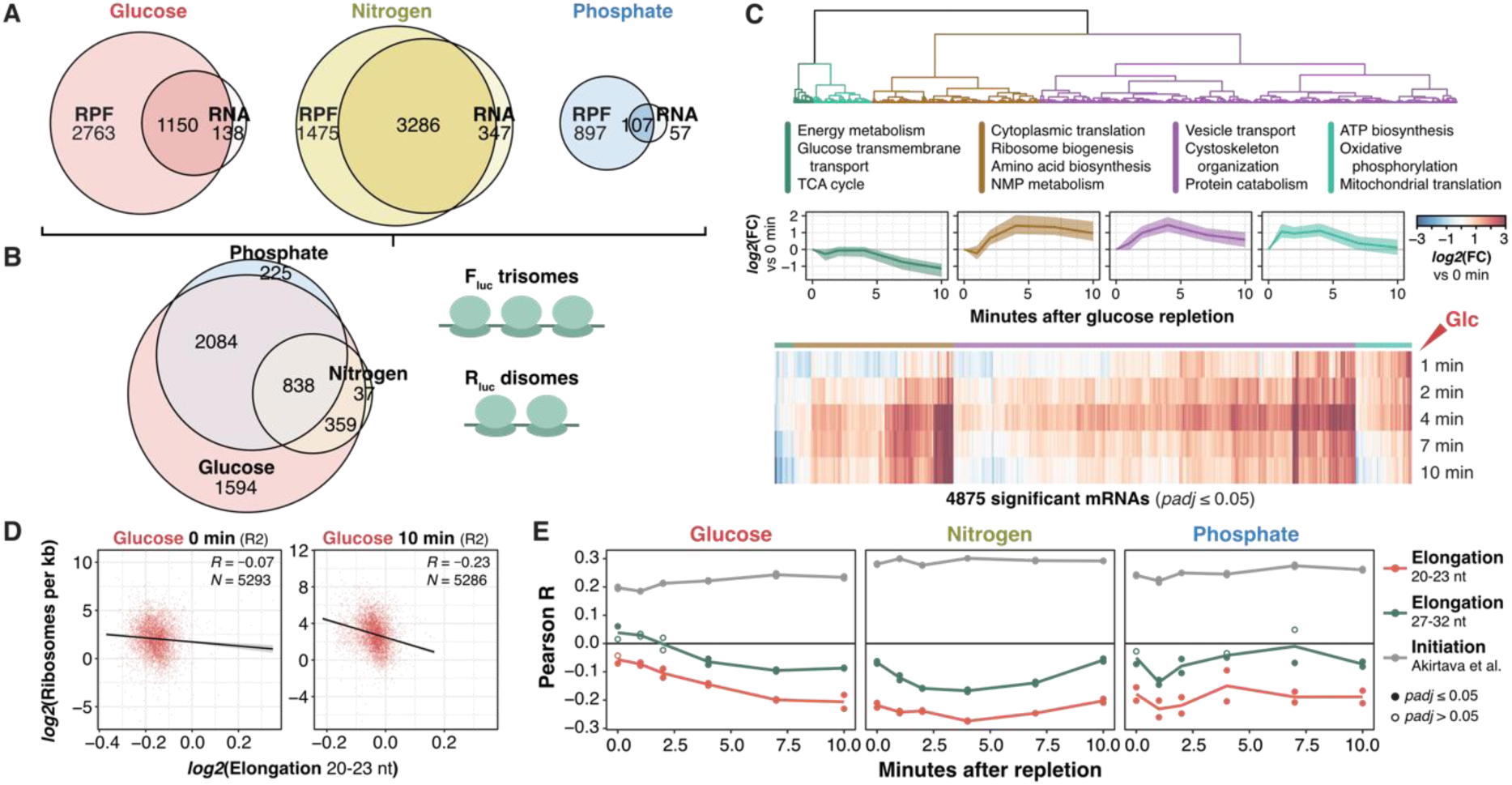
Nutrient repletion alters mRNA-specific ribosome occupancy and its relationship to codon decoding. (A) Venn diagrams of differentially expressed transcripts over the 10-min time-course in either ribosome footprints and RNA-seq, for each of the three nutrient repletions. Significant transcripts identified using *limma-voom* with a cubic spline (n_RPF_=12 and n_RNA_=6 for each nutrient repletion, *p_adj_* ≤ 0.05) (B) Right: calibrated ribosomes per transcript of the *Fluc* trisomes and *Rluc* disomes; lines show the average of n=12 per nutrient repletion. Left: Venn diagram of transcripts with differentially changing ribosomes per transcript over the 10-min nutrient repletion. Significant transcripts identified using *limma* with a cubic spline (n=12, *p_adj_* ≤ 0.05) (C) Clustering analysis of transcripts with differentially changing ribosomes per transcript upon glucose repletion. Significant transcripts were clustered by time pattern using 1 − *R_Pearson_* as distance metric and Ward’s minimum variance linkage method. For each cluster, the line plot depicts the median and the shaded IQR, as well as the top hits of the corresponding enrichment analysis of GO Biological Processes (Fisher’s exact test, *p_adj_* ≤ 0.05). Heatmap shows the average of n=2 biological replicates. (D) Number of ribosomes per kilobase of each transcript against elongation time estimate based on the geometric mean of ribosome occupancies per codon at 0 (left) and 10 min (right) upon glucose repletion (see Methods). (E) Changes in elongation time estimates over the repletion time-course. Initiation estimates calculated during normal growth from ref. ^41^. Empty dots show non-significant (*p_adj_*>0.05) correlations after Bonferroni correction. Lines show the average of n=2 biological replicates.

These relative measurements show how ribosomes are redistributed among transcripts, but not how many ribosomes each transcript carries. To measure ribosome load directly, we converted RPF and mRNA counts into calibrated values using spike-in firefly luciferase (F_luc_) and *Renilla* luciferase (R_luc_) reporter mRNAs carrying defined numbers of ribosomes (F_luc_ trisomes and R_luc_ disomes)^23^ (**Fig. 2B, right**). Ribosome load per transcript was highly correlated across biological replicates (**Extended Data Fig. 3A**) and changed for 4,875 transcripts after glucose repletion, 1,294 after nitrogen repletion, and 3,207 after phosphate repletion (**Fig. 2B, left** and **Supplementary Table 2**). The glucose response spanned most quantified transcripts, consistent with the global recovery of polysomes (**Fig. 1E, F**). In contrast, the phosphate response occurred with little change in bulk polysomes (**Fig. 1F**), consistent with a redistribution of ribosomes among transcripts.

Clustering transcripts by time revealed both nutrient-specific and shared programs. After glucose repletion, transcripts encoding cytoplasmic translation and ribosome biogenesis factors gained ribosomes; transcripts encoding vesicle transport, protein catabolism, and oxidative phosphorylation factors transiently gained ribosomes, and transcripts encoding proteins involved in energy metabolism, glucose transport, and the TCA cycle lost ribosomes (**Fig. 2C**). Transcripts encoding for cytoplasmic translation and ribosome biogenesis factors also gained ribosomes after nitrogen repletion and, by 10 min, after phosphate repletion (**Extended Data Fig. 3B, C**). Within one minute of phosphate repletion, the response was enriched for transcripts involved in membrane lipid metabolism, consistent with known links between phosphate signaling and lipid metabolism^5^.

**Extended Data Fig. 3.**
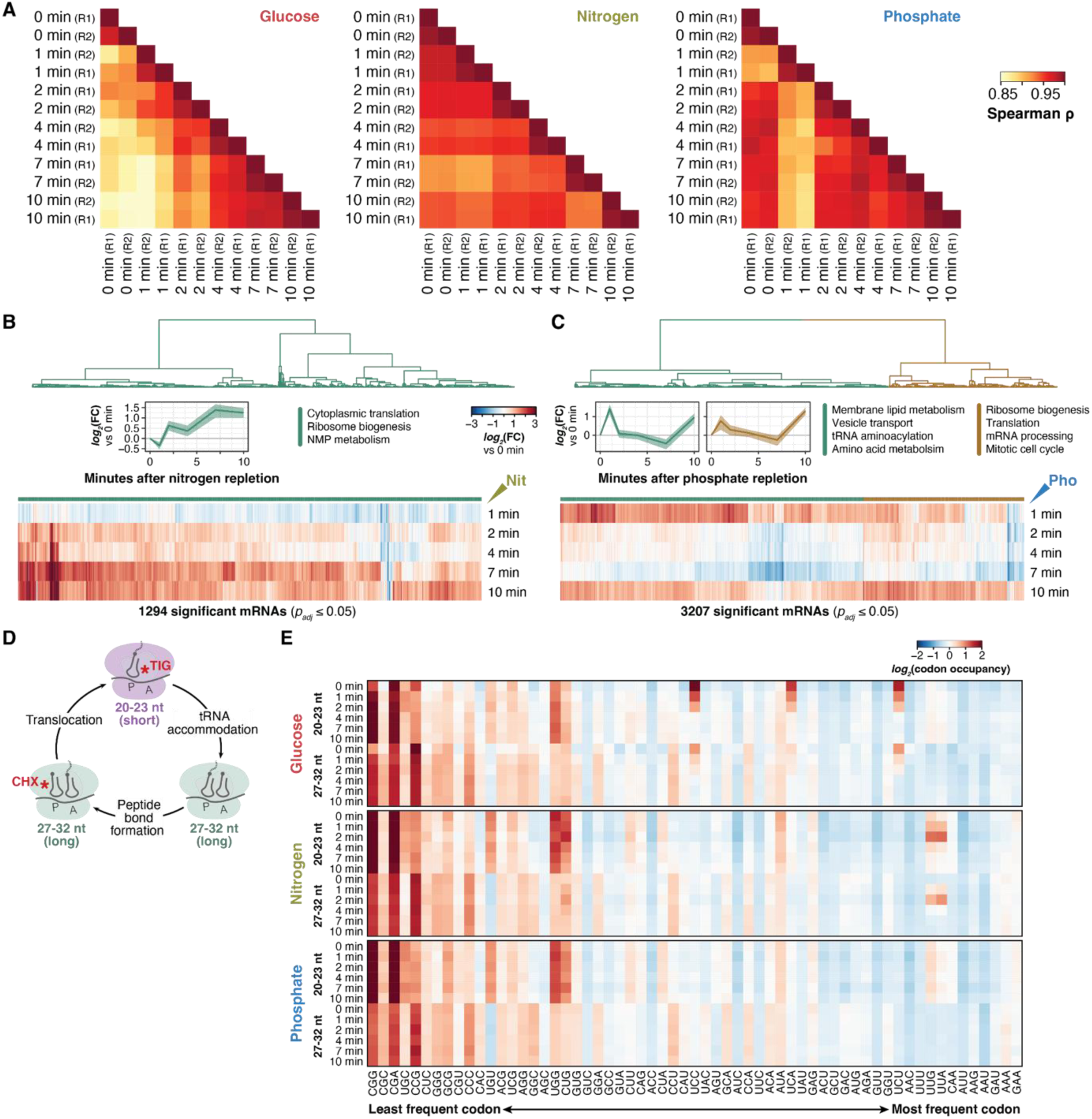
Nutrient repletion alters mRNA-specific ribosome occupancy and its relationship to codon decoding. (A) Correlation matrix among all samples for each nutrient repletion. (B-C) Clustering analysis of transcripts with differentially changing ribosomes per transcript upon nitrogen and phosphate repletions. Significant transcripts identified using *limma* with a cubic spline (n=12 for each nutrient repletion, *p_adj_* ≤ 0.05). Transcripts were clustered by time pattern using 1 − *R_Pearson_* as distance metric and Ward’s minimum variance linkage method. For each cluster, the line plot depicts the median and the shaded IQR, as well as the top hits of the corresponding enrichment analysis of GO Biological Processes (Fisher’s exact test, *p_adj_* ≤ 0.05). Heatmap shows the average of n=2 biological replicates. (D) The combination of cycloheximide and tigecycline during lysis allows the discrimination of long and short footprints during ribosome profiling, corresponding to tRNA-occupied and vacant A-site states of the ribosome (ref. ^24,25^). (E) A-site ribosome occupancies of all amino-acid-encoding codons, ordered from the least to the most frequent, averaged among all yeast coding sequences. The average of n=2 biological replicates is shown for each nutrient repletion and time point.

Ribosome load on an mRNA reflects both initiation and elongation: at a given initiation rate, slower elongation should increase the number of ribosomes per transcript. We therefore asked whether differences in elongation speed are associated with differences in ribosome load calculated after spike-in calibration. To estimate elongation speed, we used high-resolution ribosome profiling to measure codon-specific A-site occupancy, which reflects how long ribosomes pause at each codon. Adding tigecycline together with cycloheximide during lysis preserves two ribosome states that yield two classes of RPF^24,25^: long footprints (27-32 nt) corresponding to tRNA-occupied ribosomes before translocation and short footprints (20-23 nt) corresponding to ribosomes with a vacant A site after translocation, awaiting aminoacyl-tRNA (**Extended Data Fig. 3D**). These footprint classes allowed us to estimate codon-specific A-site occupancies for each nutrient and time point. Consistent with previous observations, rare codons exhibited higher ribosome occupancy than common codons, particularly among short footprints (**Extended Data Fig. 3E**). Serine and leucine codons showed transient and repletion-specific patterns. Because short footprints capture ribosomes waiting for aminoacyl-tRNA, their codon-specific occupancy provides a more direct readout of decoding time. We estimated the expected elongation time of each transcript as the geometric mean of the condition-specific A-site occupancies of its codons and compared it with spike-in-calibrated ribosome load (ref. ^42^, and *Methods*).

Ribosome load was effectively uncorrelated with estimated elongation time in glucose-deprived cells (R = −0.07) but became modestly negatively correlated after 10 min of glucose repletion (R = −0.23; **Fig. 2D, E**). Thus, transcripts enriched in codons associated with slower decoding tended to carry fewer ribosomes per mRNA, opposite to the positive relationship expected from longer ribosome residence times alone. This could reflect covariation between codon composition and other determinants of ribosome loading; for instance, rapidly decoded codons are enriched in highly expressed yeast genes, which also tend to initiate efficiently^32,34–36^. Moreover, because elongation time estimates and ribosome load are derived from the same ribosome profiling measurements, these associations should be interpreted cautiously. The negative correlation was already present in nitrogen– and phosphate-deprived cells and remained similar throughout repletion (**Fig. 2E**). This suggests that, as translation resumes after glucose deprivation, the association between ribosome load and codon composition becomes more pronounced.

We also compared ribosome load with independent estimates of translation initiation efficiency obtained for each native 5′ UTR in a massively parallel reporter assay in exponentially growing cells^43^. Because these estimates are fixed for each mRNA, they cannot capture condition-specific changes in initiation; nonetheless, they correlated positively and stably with ribosome load throughout recovery (R ∼ 0.2–0.3; **Fig. 2E**). Although these comparisons do not directly resolve the contributions of initiation and elongation, they suggest that both 5′ UTR-encoded initiation potential and codon composition are associated with transcript-specific ribosome load, with transcripts rich in rapidly decoded codons carrying more ribosomes, and that the association with codon composition depends on metabolic state, being weakest in glucose-deprived cells.

### RiboNet jointly predicts ribosome load and distribution from mRNA sequence

Ribosome load correlated with both elongation– and initiation-related sequence features, but whether transcript-specific variation in ribosome number and position can be predicted from mRNA sequence, and which regions of the transcript carry that information, remained unclear. To address these questions, we developed RiboNet, a sequence-to-function deep learning model that takes the full mature transcript as input and jointly predicts ribosome load and distribution along the transcript at codon resolution, so that sequence determinants anywhere in the transcript can be identified without predefining them (**Fig. 3A**; *Methods*). RiboNet embeds the 5′ and 3′ UTRs with a DNA language model pre-trained on fungal genomes (SpeciesLM; ref. ^44^), encodes the coding sequence (CDS) with convolutional layers, and integrates all three regions with a transformer layer (**Fig. 3A**). Its multi-task architecture generates separate predictions for each time point and footprint-length class, allowing sequence information to be learned jointly across the respective measurements (**Fig. 3A, B**). We trained independent models for glucose, nitrogen, and phosphate repletion to examine sequence-dependent regulation across distinct metabolic conditions.

**Fig. 3.**
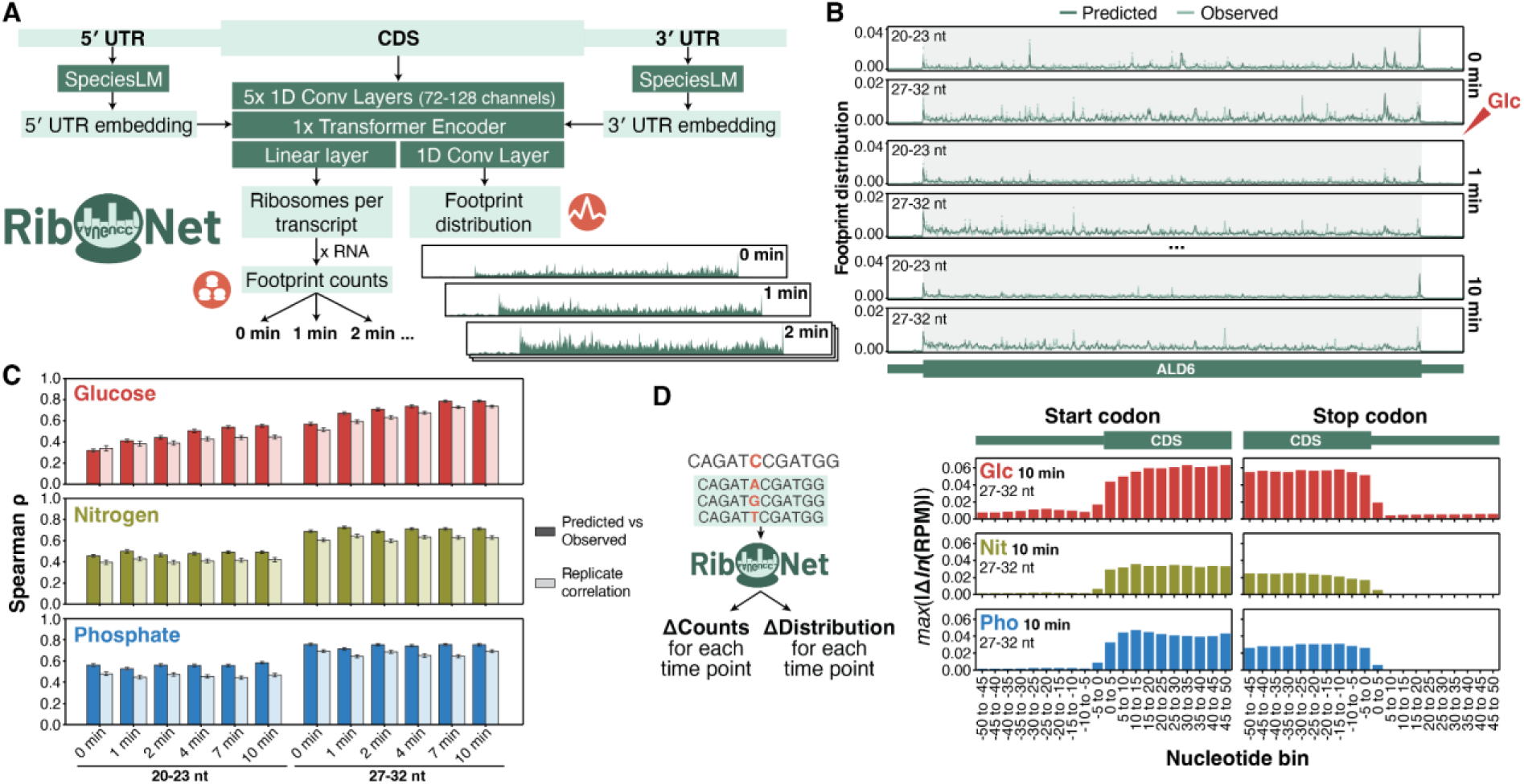
RiboNet jointly predicts ribosome load and distribution from mRNA sequence. (A) The deep learning model RiboNet takes the full mature transcript sequence as input and predicts both footprint distribution and counts simultaneously for each time point and for short or long footprints. The UTR sequences are processed through the frozen SpeciesLM model pre-trained on fungal genomes^44^. (B) Predicted and observed footprint distribution along the ALD6 transcript, measured as the fraction of footprints per position. Observed lines show the average of n=2 biological replicates. (C) Spearman correlation of each time point and long/short footprints for each of the nutrient repletion models. Bars and error bars show the average ± standard deviation among 50 random train-test splits. (D) Attribution score by *in silico* mutagenesis of every position around the start and end of the coding sequence, averaged across all transcripts. The 10-min time point is shown as an example of each nutrient repletion.

We evaluated RiboNet’s predictive performance on transcripts excluded from training, across 50 random training/test splits. Across nutrient conditions and time points, RiboNet explained ∼50-70% of the variance in footprint counts across transcripts, compared with ∼40-50% for mRNA abundance alone, with correlations across genes reaching 0.8 (**Extended Data Fig. 4A** and **Supplementary Table 3**). Removing the UTRs did not measurably reduce performance, and a linear model based on codon frequencies performed comparably to, or slightly better than, RiboNet (**Extended Data Fig. 4A)**. Sequence-based models also captured temporal changes after glucose repletion that mRNA abundance did not, whereas after nitrogen repletion mRNA abundance already accounted for much of the change (**Extended Data Fig. 4A, bottom row**). RiboNet also accurately reproduced the positional distribution of ribosome footprints along coding sequences, including differences between short and long footprints and their changes during recovery (**Fig. 3B**). Across the three nutrient conditions, prediction-observation rank correlations were generally comparable to, and occasionally exceeded, correlations between biological replicates (**Fig. 3C**, **Extended Data Fig. 4B-C** and **Supplementary Table 3**). Prediction accuracy was lower for short footprints during glucose deprivation, consistent with the reduced reproducibility of these measurements when translation was strongly suppressed (**Fig. 3C** and **Fig. 1F**).

To identify the sequence features underlying RiboNet’s predictions, we systematically substituted individual nucleotides *in silico* and quantified their effects on predicted ribosome load and distribution (**Fig. 3D**). This analysis revealed that the CDS contained most of the sequence information influencing predicted ribosome load per mRNA, with attribution rising at the start codon and falling at the stop codon (**Fig. 3D**). Importantly, predictive information was also detected in untranslated regions, particularly in the 5′ UTR during glucose repletion, although removing the UTRs did not reduce average predictive performance (**Extended Data Fig. 4A**), suggesting that UTR features act on a subset of transcripts or overlap with information in the CDS. Taken together, these analyses establish that RiboNet reproduces codon-resolution ribosome distributions and provide a basis for identifying sequence determinants of ribosome load and distribution along transcripts across changing conditions.

**Extended Data Fig. 4.**
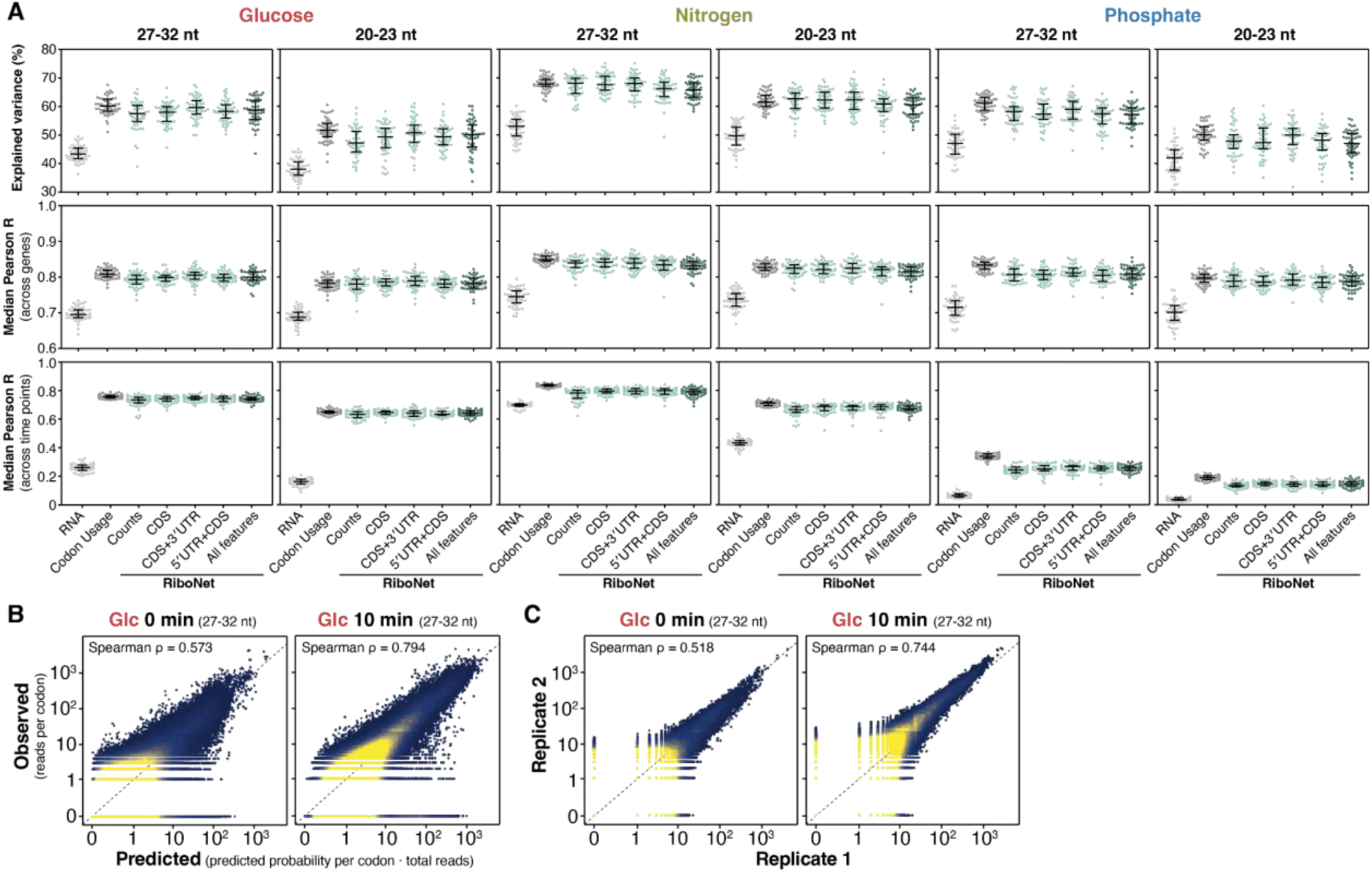
RiboNet jointly predicts ribosome load and distribution from mRNA sequence. (A) Benchmarking of the full RiboNet, ablated versions (removing 5′ UTR, 3′ UTR, both, or training on ribosome counts-only), and the baselines of RNA abundance and a multiple linear regression of gene codon frequencies. Three metrics are computed on the held-out test set (percentage of explained variance across genes and time points, median Pearson correlation coefficient across genes, and median Pearson correlation coefficient across time points) for each of the three repletions, separating between short and long RPFs. 50 iterations of each model were trained on different train-test splits. The central line and error bars correspond to the median and IQR. (B) Scatter plot of predicted and observed ribosome counts per position (i.e. footprint distribution) for all genes in the test set with the corresponding Spearman correlation coefficient. (C) Scatter plot and correlation of observed ribosome counts per position between n=2 biological replicates.

### Nutrient-dependent pausing at specific codons is linked to changes in tRNA charging

The elongation-time estimates above treat each codon’s dwell time as independent of its sequence context. To determine whether nutrient-dependent changes in ribosome occupancy depend primarily on the A-site codon or also on the neighboring sequence, we used RiboNet to identify their sequence determinants. For each repletion, we selected the 5,000 positions (P-site codons in individual transcripts) with the largest observed changes in ribosome occupancy across time points. At each position, we mutated every nucleotide in a window spanning the ribosome *in silico* and asked how each substitution altered the model’s predicted change in occupancy. This analysis revealed distinct codon-specific responses to glucose and nitrogen repletion, in each case concentrated on the A-site codon, consistent with measured ribosome occupancies (**Extended Data Fig. 3E**). RiboNet attributed the decrease in occupancy after glucose repletion to serine UCU and UCC codons in the A site, particularly in short footprints corresponding to ribosomes with vacant A sites (**Fig. 4A** and **Extended Data Fig. 5A**). In contrast, it attributed the transient increase in occupancy after nitrogen repletion, most prominent during the first two minutes, to leucine UUA and UUG codons in the A site (**Fig. 4B** and **Extended Data Fig. 5B**). No comparable codon-specific responses were detected following phosphate repletion (**Extended Data Fig. 3E**). Thus, these nutrient-dependent occupancy changes are associated primarily with the identity of the A-site codon, consistent with changes in the decoding step.

**Fig. 4.**
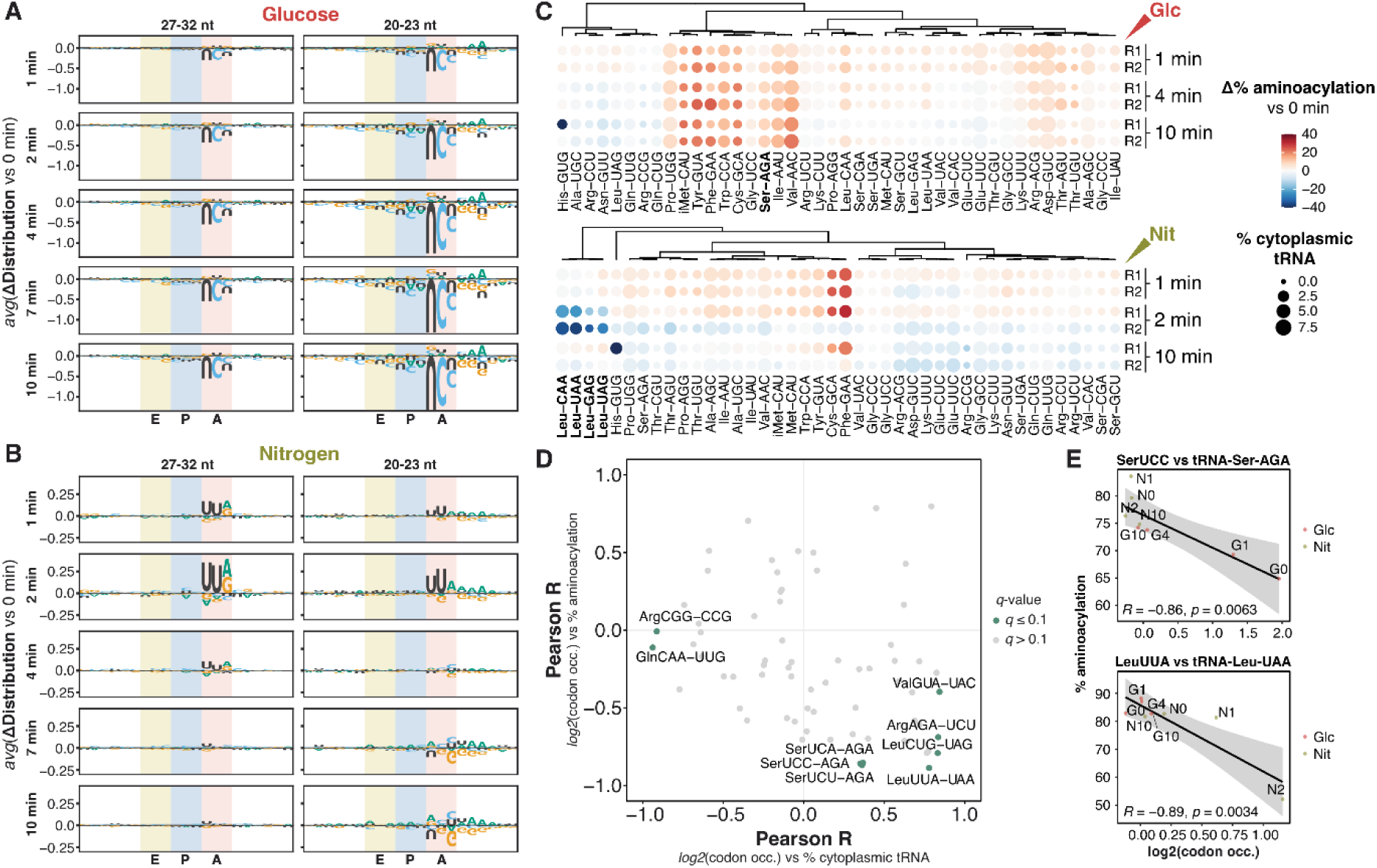
Nutrient-dependent pausing at specific codons is linked to changes in tRNA charging. (A-B) Sequence determinants of changes in ribosome occupancy after glucose (A) and nitrogen (B) repletion, identified by *in silico* mutagenesis with RiboNet. For the 5,000 sites whose observed occupancy changed most across time points, each nucleotide in a window around the ribosome was mutated *in silico*, and its effect on the predicted change in footprint probability at that site (time point vs 0 min) was computed. Sites are aligned on the P-site codon; shaded columns mark the E, P, and A sites. Letter height shows the average contribution of each nucleotide to the change; letters below zero mark nucleotides whose contribution to occupancy decreased relative to 0 min. Left, long (27-32 nt) footprints; right, short (20-23 nt) footprints. (C) Abundance and change in aminoacylation of all cytoplasmic tRNA anticodon families upon glucose and nitrogen repletion (n=2 biological replicates per time point). (D) Pearson correlation of all codon-anticodon pairs^45^ across glucose and nitrogen repletion. Codon occupancies of short RPFs were correlated both against cognate tRNA abundance (x axis) and tRNA aminoacylation (y axis). Codons where correlations have a *q*-value ≤ 0.1 are labeled. (E) Scatter plots of two selected correlations in D.

To investigate whether these changes were associated with the availability of cognate charged tRNAs, we measured tRNA abundance and charged fractions at selected time points during glucose and nitrogen repletion using mim-tRNAseq^46^. We did not detect significant changes in the abundance of individual tRNAs during either repletion, whereas the fraction of aminoacylated tRNAs changed across several anticodon families (**Fig. 4C** and **Supplementary Table 4**). In glucose-deprived cells, tRNA-Ser-AGA, the most abundant serine isoacceptor, which decodes UCU and UCC, was markedly undercharged, and its aminoacylation increased to ∼75% within 4 min of glucose repletion (**Extended Data Fig. 5C**). Other serine tRNAs also exhibited increased charging, but from a higher baseline (**Fig. 4C**). Nitrogen repletion predominantly affected leucine tRNAs: all four leucine isoacceptors, including tRNA-Leu-CAA and tRNA-Leu-UAA, which decode UUG and UUA, respectively, transiently lost charging at 2 min and recovered by 10 min (**Fig. 4C** and **Extended Data Fig. 5C**). Changes in tRNA aminoacylation were only partially reflected in the intracellular abundance of the corresponding amino acids (**Extended Data Fig. 5D-E** and **Supplementary Table 4**): leucine fell transiently 2-4 min after nitrogen repletion, coinciding with the loss of leucine charging, whereas serine was similarly low in glucose– and nitrogen-deprived cells, although tRNA-Ser-AGA was undercharged only in glucose-deprived cells.

We next examined the relationship between codon-specific ribosome occupancy and cognate tRNA abundance or aminoacylation across the eight sampled conditions and time points. Several codons exhibited negative correlations between short-footprint occupancy and the fraction of aminoacylated cognate tRNAs (**Fig. 4D**). These included the serine and leucine codons identified by RiboNet, alongside additional codons (**Fig. 4E**). The strongest examples included Ser UCC and Leu UUA, whose measured occupancies were inversely correlated with the aminoacylation of their cognate tRNAs (R = −0.86 and −0.89, respectively; **Fig. 4E**). Ser UCC occupancy, for example, was ∼4-fold above the transcript average in glucose-deprived cells, when tRNA-Ser-AGA charging was lowest, and returned to average within 10 min (**Fig. 4E**). Similar associations were observed for long footprints (**Extended Data Fig. 5F-G**). In contrast, occupancy at few codons correlated with cognate tRNA abundance, and these differed between short and long footprints (**Fig. 4D** and **Extended Data Fig. 5F, G**). Although several correlations were suggestive at a false discovery rate of 10%, none remained significant at 5%, limiting the strength of the conclusions drawn from these associations. Nonetheless, the codons with the strongest associations were the same serine and leucine codons highlighted by RiboNet, and their occupancy changes matched the timing of cognate tRNA charging.

To examine which transcripts might be particularly sensitive to changes in decoding at these codons, we assessed their enrichment across functional gene categories. Ser UCU and UCC were enriched in transcripts involved in cell wall organization and cytoplasmic translation, whereas Leu UUG was associated with cytoplasmic translation and Leu UUA with respiratory electron transport (**Extended Data Fig. 5H**). These patterns indicate that the codons exhibiting nutrient-dependent occupancy changes are unevenly distributed across functionally related transcripts, although the enrichment for cytoplasmic translation partly reflects the preference for these codons in highly expressed genes^47^.

Taken together, these data link nutrient-dependent ribosome pausing at selected codons with changes in cognate tRNA charging and suggest that the regulation of aminoacylation during metabolic recovery involves factors beyond amino acid availability.

**Extended Data Fig. 5.**
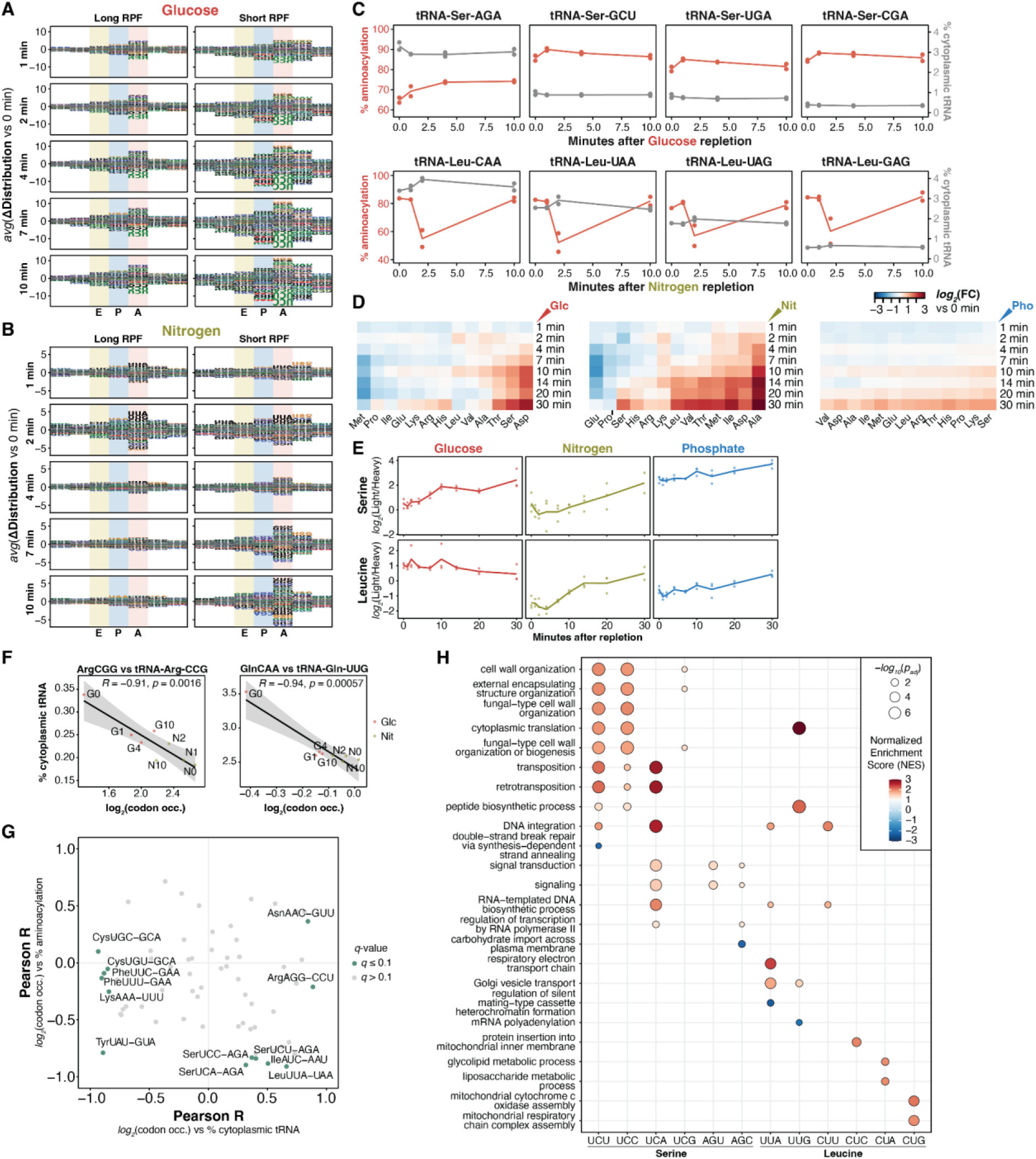
Nutrient-dependent pausing at specific codons is linked to changes in tRNA charging. (A-B) As in Fig. 4A, B, but averaging per codon; letter height shows the average effect of each codon substitutions. (C) Abundance and aminoacylation of all serine and leucine tRNAs across the time course of glucose and nitrogen repletions, respectively. Source data corresponds to Fig. 4C. Lines show the average of n=2 biological replicates. (D) Targeted metabolomics of all detected amino acid internal standards across the 30-minute time course for each nutrient repletion. (E) Serine and leucine metabolite abundance, same source data as D. Heatmap in D and lines in E show the average of n=3 biological replicates. (F) Scatter plots of two selected correlations in Fig. 4D. (G) Same as Fig. 4D, but using codon occupancies of long RPFs. (H) Gene Set Enrichment Analysis (GSEA) of all genes ordered by codon frequency, indicating the top 2 most significant GO Biological Processes (gene-label permutation test, *p_adj_* ≤ 0.05) per codon and Normalized Enrichment Score sign.

### Nutrient repletion induces rapid structural changes in the translation machinery

To investigate how the translation machinery responds to metabolic changes, we monitored protein structure during nutrient repletion using limited proteolysis coupled to mass spectrometry (LiP-MS, ref. ^48,49^). This approach detects changes in local protein structure through differential accessibility to proteolytic cleavage (**Fig. 5A**). Accessibility was normalized to protein abundance measured in trypsin-only controls, so that structural changes could be distinguished from changes in protein levels (see *Methods*). Across the first 20 minutes of nutrient repletion, substantially more proteins exhibited changes in proteolytic accessibility than in abundance (**Fig. 5B** and **Supplementary Table 5**). These structural dynamics occurred rapidly and differed markedly between nutrients, with 3,204 significantly changing LiP-MS peptides following glucose repletion, compared with 1,782 following phosphate and 1,112 following nitrogen repletion (**Extended Data Fig. 6A-C**). Among proteins involved in translation, up to 60% exhibited changes in proteolytic accessibility, similar to the proportion among all quantified proteins (**Extended Data Fig. 6D** and **Fig. 5B**). After glucose repletion, these changes peaked within 1-2 min, whereas changes in the abundance of translation-related proteins developed over 10-20 min (**Fig. 5C**).

**Fig. 5.**
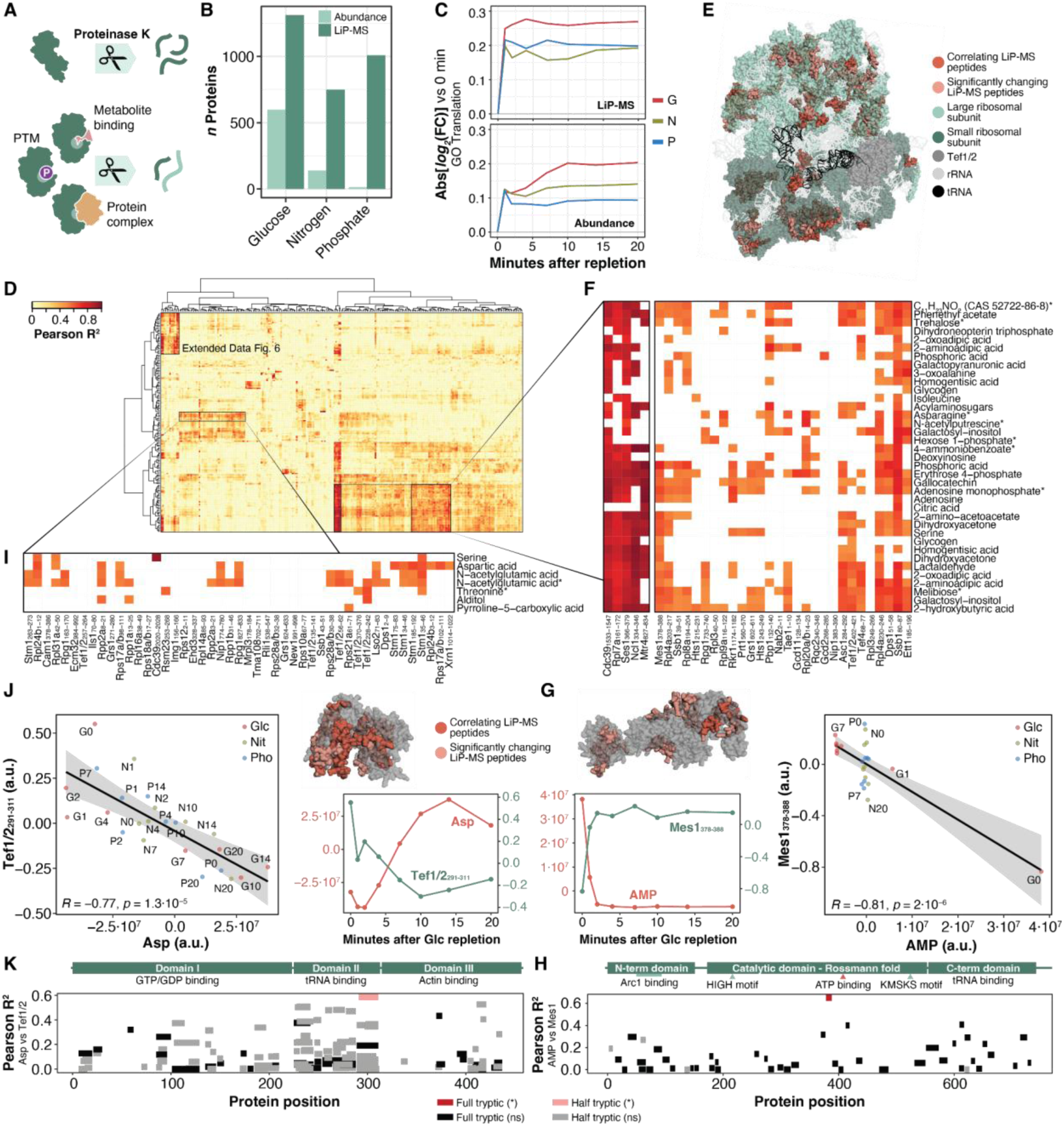
Nutrient repletion induces rapid structural changes in the translation machinery. (A) Limited Proteolysis coupled to MS detects protein conformational changes caused by PTMs or molecular interactions. (B) Number of proteins either significantly changing their abundance or with at least one peptide differentially changing structure, for each of the 20-minute nutrient repletions. Significant features identified using *limma* with a cubic spline (n=24 for each nutrient repletion, *p_adj_* ≤ 0.05). (C) Median absolute change of all detected proteins (abundance) and peptides (LiP-MS) associated with translation based on GO Biological Process annotation, colored for each nutrient repletion. (D) Correlation matrix of significantly changing translation-related peptides with significantly changing identified metabolites. Only peptides and metabolites with at least one significant correlation are shown (*p_adj_* ≤ 0.05, local FDR correction). Hierarchical clustering uses the Euclidean distance metric and Ward’s minimum variance linkage method. (E) Ribosome structure (PDB 8Z70) that highlights all significantly changing peptides in any of the three nutrient repletions and whether they correlate significantly with any metabolite in D. (F) Augmented section of one cluster of the correlation matrix, coloring only correlations with an FDR-corrected *p*-value < 0.05. Metabolites marked with an asterisk indicate MS2-based metabolite annotation. (G) Median time-course pattern of Mes1_378-388_ peptide and AMP across the glucose repletion (n=3 per time point), protein structure (AlphaFold EBI), and scatter plot of the measured correlation. (H) Woods’ plot of the correlation of all Mes1 peptides with AMP. (I) Augmented section of a second cluster of the correlation matrix. (J) Median time-course pattern of Tef1/2_257-264_ peptide and aspartate across the glucose repletion (n=3 per time point), protein structure (PDB 1F60), and scatter plot of the measured correlation. (K) Woods’ plot of the correlation of all Tef1/2 peptides with aspartate.

To identify metabolic changes associated with these structural responses, we integrated matched LiP-MS and metabolomic measurements across the three nutrient-repletion experiments. Correlation analysis of significantly changing peptides and metabolites revealed distinct groups of covarying signals among translation-related proteins (**Fig. 5D** and **Supplementary Table 5**). Strongly correlated peptides mapped to several aminoacyl-tRNA synthetases (Ses1, Mes1, Dps1), the elongation factor eEF1A (Tef1/2), and proteins from both ribosomal subunits (**Fig. 5E** and **Extended Data Fig. 6E**). The associated metabolites include energy-related compounds, such as AMP and sugar phosphates, as well as amino acid derivatives (**Fig. 5D** and **Extended Data Fig. 6F**).

The largest cluster contained more than half of the metabolite-correlated peptides and included many energy-related metabolites, such as AMP, adenosine, and sugar phosphates (**Fig. 5F**). This cluster included peptides from 14 of the 20 cytosolic aminoacyl-tRNA synthetases with at least one significantly changing peptide. Methionyl-tRNA synthetase (Mes1) provided a particularly informative example: among several peptides exhibiting altered proteolytic accessibility, the peptide spanning residues 378-388 was strongly anti-correlated with AMP (*R* = −0.81, *P* = 2 × 10⁻^6^; **Fig. 5G**). Its accessibility changed within one minute of glucose repletion, coinciding with the sharp decline in cellular AMP. This peptide maps to the ATP-binding region of Mes1 (**Fig. 5H**), suggesting a potential connection between adenylate metabolism and structural changes near the enzyme’s catalytic site. Similar responses were observed in other aminoacyl-tRNA synthetases, including the seryl-tRNA synthetase Ses1, whose ATP-binding region contained a peptide strongly anti-correlated with AMP (*R* = −0.91; **Extended Data Fig. 6G-H**). Because these AMP correlations rest largely on the difference between glucose-deprived cells and later time points (**Fig. 5G** and **Extended Data Fig. 6H**), and because aminoacylation consumes ATP, these changes may reflect the recovery of nucleotide binding at the active site as energy status is restored, consistent with the increased charging of many tRNAs after glucose repletion (**Fig. 4C**). For serine, this coincided with increased charging of tRNA-Ser-AGA and reduced pausing at UCU and UCC codons (**Fig. 4A, C**), suggesting that energy-dependent changes in synthetase activity may contribute to altered decoding.

Two additional clusters involved metabolites that changed after nitrogen repletion. The first contained peptides correlating with amino acids and their derivatives, including serine and aspartate, across glucose and nitrogen repletion (**Fig. 5I**). This group included the ribosome preservation factor Stm1 and multiple peptides from the elongation factor eEF1A Tef1/2 (**Fig. 5J**). A Tef1/2 peptide in domain II, which binds aminoacyl-tRNA, exhibited a strong negative correlation with aspartate (*R* = −0.77, *P* = 1.3 × 10⁻^5^; **Fig. 5J**). It was the only Tef1/2 peptide significantly correlated with aspartate (**Fig. 5K**). The second cluster was dominated by metabolites responding to nitrogen repletion, including tricarboxylic acid cycle intermediates and nucleosides (**Extended Data Fig. 6I**). Leucyl-tRNA synthetase showed metabolite-associated structural changes, including a positive correlation between guanosine and the accessibility of a peptide in its C-terminal tRNA-binding domain (*R* = 0.75, P = 2.5 × 10⁻⁵; **Extended Data Fig. 6G-H**). These changes occurred alongside the transient loss of leucine tRNA charging and increased ribosome pausing at UUA and UUG leucine codons following nitrogen repletion (**Fig. 4B-C**).

Taken together, these findings identify metabolite-associated structural changes in aminoacyl-tRNA synthetases that coincide with nutrient-specific alterations in cognate tRNA charging and ribosome pausing. They are consistent with a model in which metabolic regulation of tRNA aminoacylation contributes to rapid changes in codon decoding during translational recovery.

**Extended Data Fig. 6.**
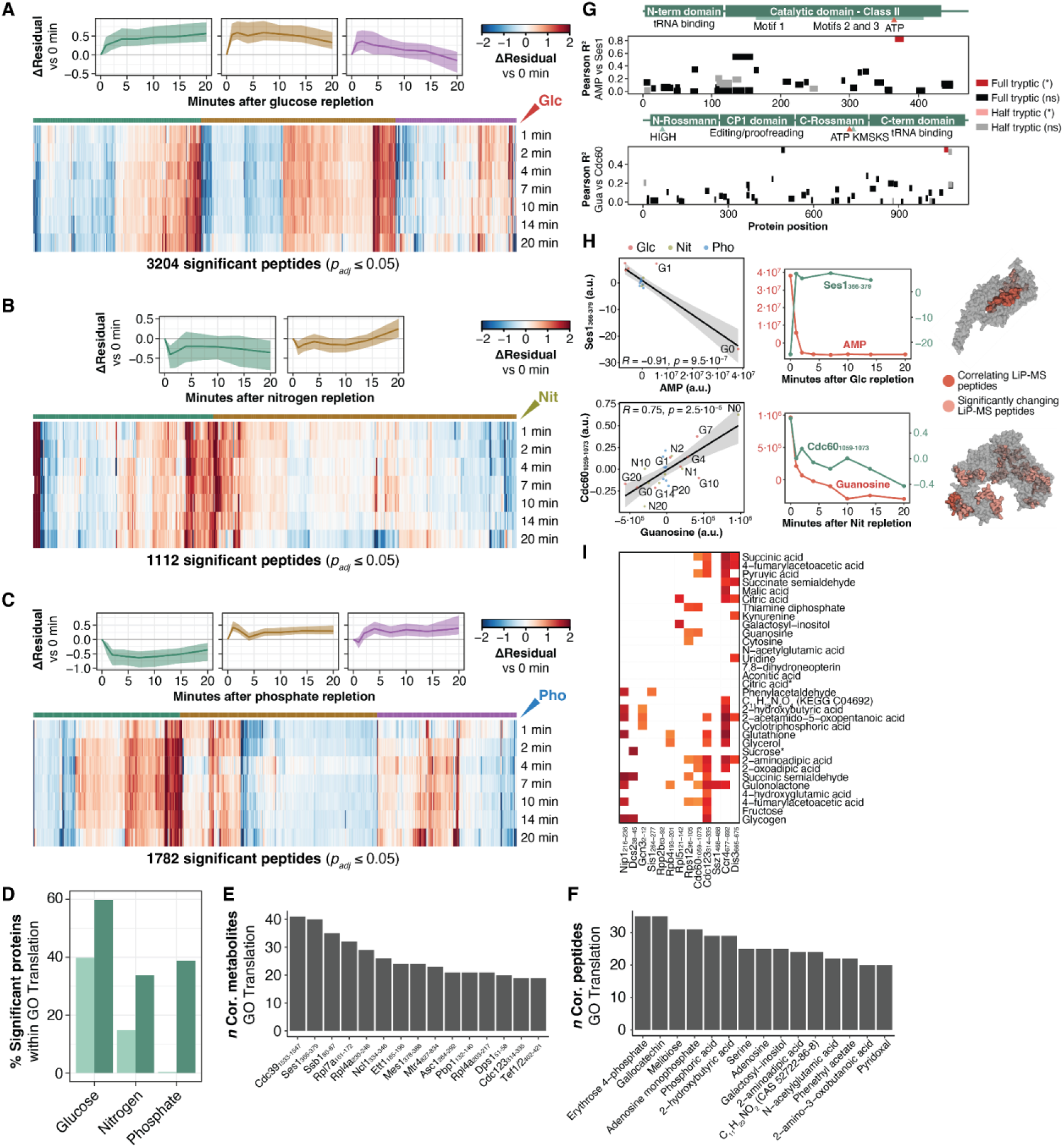
Nutrient repletion induces rapid structural changes in the translation machinery. (A-C) Clustering analysis of all LiP-MS significantly changing peptides across the three nutrient repletions. Significant peptides identified using *limma* with a cubic spline (n=24 for each nutrient repletion, *p_adj_* ≤ 0.05). Peptides were clustered by time pattern using 1 − *abs*(*R_Pearson_*) as the distance metric and Ward’s minimum variance linkage method. For each cluster, the line plot depicts the median and the shaded IQR. The sign of changes in PK accessibility has no meaning, and thus opposite but similar patterns can cluster together. Heatmaps show the average of n=3 biological replicates. (D) Percentage of translation-annotated proteins either significantly changing their abundance or with at least one peptide differentially changing structure, for each of the 20-minute nutrient repletions. (E-F) Number of significantly top-15 most common correlating peptides and metabolites among all translation-annotated proteins. (G) Woods’ plot of the correlation of all Ses1 and Cdc60 peptides with AMP and guanosine, respectively. (H) Median time-course pattern of two selected peptides along the nutrient repletions (n=3 per time point), scatter plot of the measured correlation, and their location on the protein structures (AlphaFold EBI). (I) Augmented section of one cluster of the correlation matrix in Fig. 5E, coloring only correlations with a *p_adj_* ≤ 0.05 (local FDR correction). Metabolites marked with an asterisk indicate MS2-based metabolite annotation.

### A conserved 5′ UTR motif, GAUY, increases protein output

Having identified sequence determinants of codon-specific ribosome positioning, we next examined regulatory elements associated with transcript-level ribosome load. We applied TF-MoDISco^50^, which groups high-attribution positions into recurrent motifs, to RiboNet attribution scores, analyzing the 5′ UTR, CDS, and 3′ UTR separately to account for differences in attribution magnitude between these regions (**Fig. 3D**). Because attributions were calculated relative to 0 min, the resulting motifs represent sequence features associated with changes in footprint counts after repletion (**Fig. 6A**). Coding regions were enriched for known inhibitory codons, including Arg CGA, Arg CGG, and Pro CCG^51,52^, as well as codon pairs and proline-rich sequences, which are known to slow peptide bond formation^53–55^. Attribution scores in 3′ UTRs were generally weaker, and predominantly associated with A/U-rich sequences. In contrast, the 5′ UTR analysis identified two candidate regulatory motifs, GAUY and AGGCUG, which did not match annotated yeast RNA motifs in databases^56,57^.

**Fig. 6.**
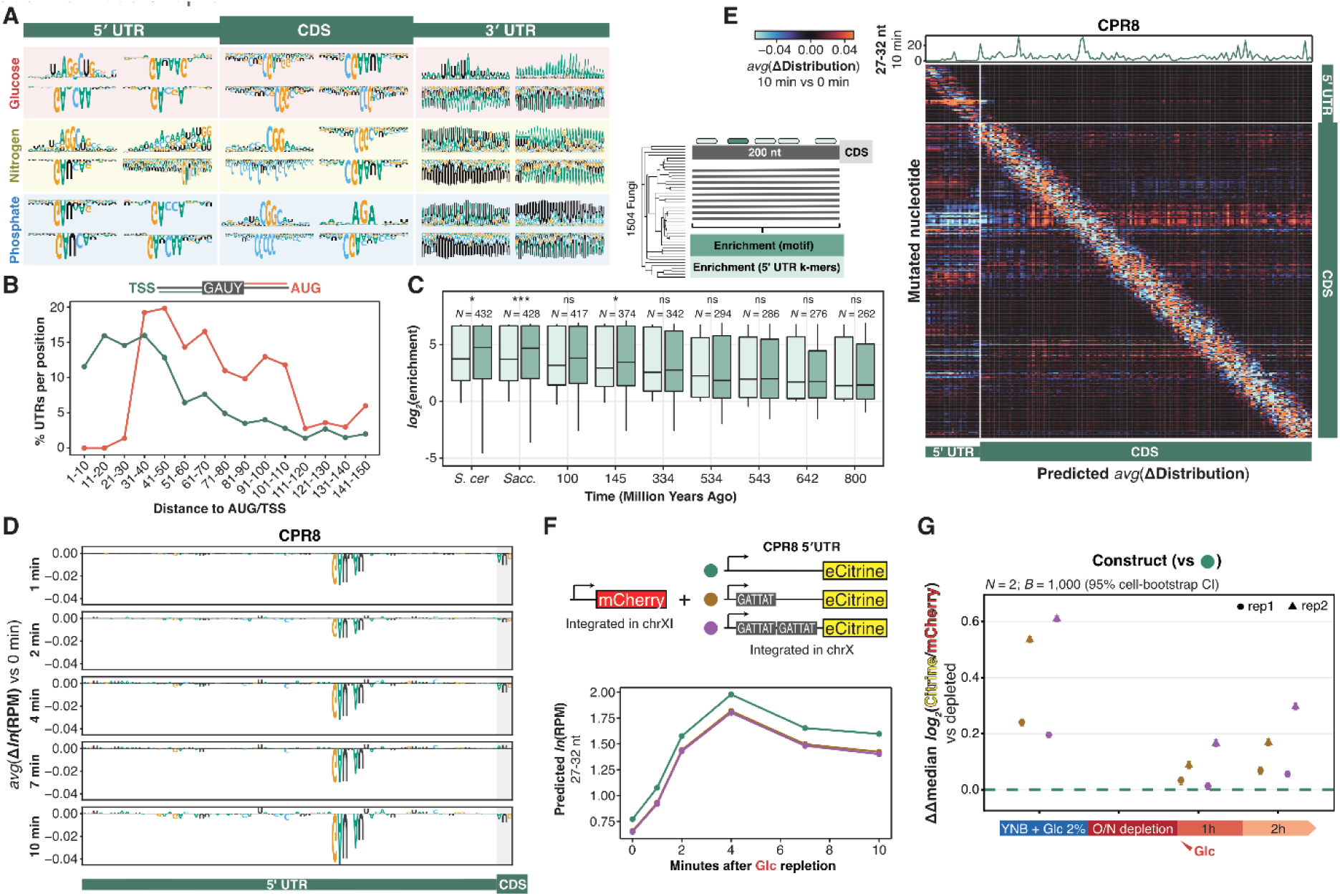
A conserved 5′ UTR motif, GAUY, increases protein output. (A) Summary of TF-MoDISco^50^-identified motifs regulating RPF counts. Attribution scores per position are determined by *in silico* mutagenesis of all genes for each region and condition relative to 0 min. (B) Location of all detected high-confidence GAUY seqlets in comparison to the start codon and the transcript start site (TSS), as a percentage of the genes with 5′ UTR length of a certain size. (C) Simple Enrichment Analysis (SEA)^59^ of the GAUY seqlet of each gene among the 200 nt upstream sequences of its homologs at each evolutionary distance. Enrichment is compared to the median background enrichment of random 5′ UTR k-mers of the same size, and significance determined by a one-sided Wilcoxon signed-rank test (ns *p*>0.05, * *p*≤0.05, ** *p*≤0.01, *** *p*≤0.001). (D) *In silico* mutagenesis of the CPR8 5′ UTR sequence, taking the average score of all 3 mutations per position and relative to 0 min. (E) Effect on RPF probability per position of the *in silico* mutagenesis of the first 500 nucleotides of the CPR8 transcript. The line plot on the top shows the observed long RPFs per position after 10 min of glucose repletion. (F) Designed synthetic sequences of CPR8 5′ UTR in front of the eCitrine reporter and their RiboNet-predicted RPMs along the glucose repletion. All reporters are integrated in chromosome X and normalized to mCherry under identical ENO2 promoter and ADH1 terminator at chromosome XI. (G) Change in median log_2_(Citrine/mCherry) relative to the glucose-depleted condition, shown as the difference between the native or duplicated GAUY construct and the scrambled control (dashed line). Points are biological replicates (n = 2). Error bars are 95% CIs from bootstrapping cells within each sample (1,000 iterations) and reflect only within-replicate sampling uncertainty.

We focused on GAUY because of its prevalence across endogenous transcripts (909 genes, versus 117 for AGGCUG). Applying a stringent attribution-score threshold identified high-confidence motif occurrences in 433 genes (**Extended Data Fig. 7A** and **Supplementary Table 6**). These sites were preferentially located 30-70 nucleotides upstream of the start codon, frequently near annotated transcript start sites (**Fig. 6B**). Comparative analysis of fungal orthologs revealed significant enrichment of GAUY at evolutionary distances extending to 145 million years, relative to random 5′ UTR k-mers of the same length (**Fig. 6C** and **Supplementary Table 6**). GAUY-containing genes showed no significant enrichment for particular biological processes or codon usage patterns (**Supplementary Table 6**).

For experimental validation, we prioritized GAUY-containing 5′ UTRs without upstream AUG codons, with motif occurrence downstream of the main annotated transcription start site^58^, and conserved among fungal orthologs (**Extended Data Fig. 7A-C**). Two candidates, *CPR8* and *TYW1*, contained GAUUAU and GAUCA sequences in their 5′ UTRs, respectively. In both transcripts, *in silico* mutagenesis assigned the strongest 5′ UTR attribution to the motif positions themselves from 1 min after glucose repletion onwards (**Fig. 6D** and **Extended Data Fig. 7D**). Substitutions in the motif region also changed the predicted distribution of footprints along the transcript, in both the 5′ UTR and the CDS (**Fig. 6E** and **Extended Data Fig. 7E**). The model therefore predicted that GAUY affects both the number and the positions of ribosomes during recovery.

To test these predictions, we generated chromosomally integrated eCitrine reporters carrying the *CPR8* or *TYW1* 5′ UTRs with zero (scrambled), one (native), or two copies of the GAUY motif. eCitrine fluorescence was normalized to an independently integrated mCherry control expressed under the same promoter and terminator. RiboNet predicted reduced CDS footprint counts for the motif-containing constructs during glucose repletion (**Fig. 6F** and **Extended Data Fig. 7F**). To confirm that the assay detects translational differences, we first tested eCitrine variants with slow and fast elongation kinetics^29^, confirming the expected differences in the eCitrine-to-mCherry ratio (**Extended Data Fig. 7G**). Immunoblotting confirmed that all strains expressed both proteins at the expected sizes (**Extended Data Fig. 7I**). Citrine-to-mCherry ratios increased after glucose repletion for all three constructs (**Extended Data Fig. 7H**). Relative to the depleted state, the increase was larger for the motif-containing constructs than for the motif-free 5′ UTR, and largest for the two-copy construct in most replicates (**Fig. 6G** and **Extended Data Fig. 7J-L**). Motif-containing constructs also showed higher eCitrine-to-mCherry ratios in cells that had never been deprived of glucose as well as 1 h and 2 h after repletion (**Extended Data Fig. 7H**), indicating that the motif affects protein output during steady-state growth, not only during recovery.

**Extended Data Fig. 7.**
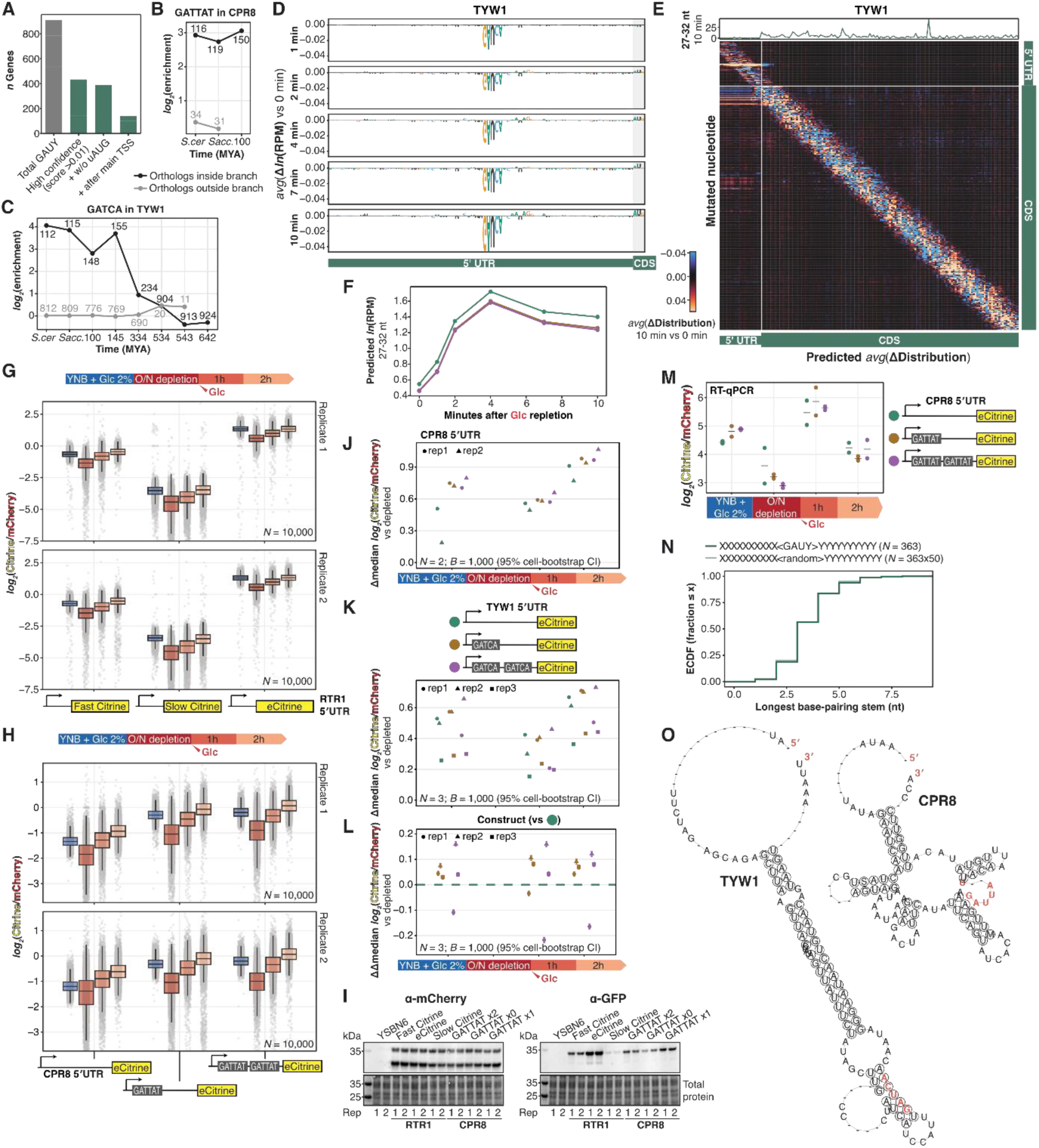
A conserved 5′ UTR motif, GAUY, increases protein output. (A) Number of GAUY seqlets and the several filters applied to subset high-confidence genes and prioritize candidate hits for experimental validation. (B) Simple Enrichment Analysis (SEA) of the GAUUAU motif among the 200 nt upstream sequences of all CPR8 orthologs. (C) Same as B, but for the GAUCA motif among TYW1 orthologs. (D) *In silico* mutagenesis of the TYW1 5′ UTR sequence, taking the average score of all 3 mutations per position and relative to 0 min. (E) Effect on RPF probability per position of the *in silico* mutagenesis of the first 500 nucleotides of the TYW1 transcript. The line plot on the top shows the observed long RPFs per position after 10 min of glucose repletion. (F) Designed synthetic sequences of TYW1 5′ UTR in front of the eCitrine reporter and their RiboNet-predicted RPMs along the glucose repletion. (G) Citrine-to-mCherry ratios for each of the synonymous citrine strains along all 4 depletion-repletion conditions analyzed. (H) Citrine-to-mCherry ratios for each of the synthetic CPR8 5′ UTR strains along all 4 depletion-repletion conditions analyzed. (I) Immunoblotting of α-mCherry and α-GFP of all the synonymous-citrine and CPR8-5′ UTR strains under the YNB+2% glucose condition, compared to the non-transformed paternal YSBN6 strain. (J) Change in median log_2_(Citrine/mCherry) relative to the glucose-depleted condition for each CPR8 5′ UTR strain. Points are biological replicates (n = 2). (K) Change in median log_2_(Citrine/mCherry) relative to the glucose-depleted condition for each TYW1 5′ UTR strain. Points are biological replicates (n = 3). (L) Change in median log_2_(Citrine/mCherry) relative to the glucose-depleted condition, shown as the difference between the native or duplicated GAUY variant and the scrambled control (dashed line). Points are biological replicates (n = 3). In J-L, error bars are 95% CIs from bootstrapping cells within each sample (1,000 iterations) and reflect only within-replicate sampling uncertainty. (M) Citrine-to-mCherry RNA ratios for each of the synthetic CPR8 5′ UTR strains determined by RT-qPCR. Points show the average of technical triplicates, lines show the average of n=2 biological replicates. (N) Empirical cumulative distribution function (ECDF) of the longest base-pairing stretch of the 10 upstream against the 10 antiparallel downstream flanking nucleotides around the GAUY motif, compared to 5′ UTR random k-mers. (O) Predicted CPR8 and TYW1 5′ UTR structures based on the simultaneous orthologs-alignment and folding by LocARNA^60^.

Reporter mRNA ratios remained comparable between constructs (**Extended Data Fig. 7M**), supporting a post-transcriptional contribution to the differences in protein output. The increased reporter fluorescence contrasted with the lower CDS footprint counts predicted by RiboNet. This discrepancy suggests that GAUY-containing sequences may affect the relationship between ribosome occupancy and protein production, potentially through changes in ribosome transit or other aspects of translation.

We next examined whether GAUY motifs were associated with conserved RNA secondary structures. High-confidence motif-containing transcripts showed no enrichment for extended local base-pairing relative to random 5′ UTR sequences (**Extended Data Fig. 7N**). Structure-guided alignment of the 5′ UTR sequences of *CPR8* and *TYW1* orthologs likewise revealed no consistent structural arrangement or substantial evidence of compensatory covariation (**Extended Data Fig. 7O**). These observations suggest that the regulatory effects of GAUY are primarily associated with its sequence rather than a conserved secondary structure.

Taken together, these findings identify GAUY as a conserved 5′ UTR element that increases protein output post-transcriptionally, both during steady-state growth and after glucose repletion.

## DISCUSSION

Restoring protein synthesis after nutrient deprivation requires cells to adjust translation to rapid changes in cellular metabolism. By combining time-resolved multi-omics measurements with sequence-based modeling, we found that glucose, nitrogen, and phosphate repletion act on different layers of gene expression and, most strikingly, that metabolic state changes the decoding of specific codons within minutes, in step with the charging of their cognate tRNAs. These findings extend current models of nutrient-dependent translational control, which have largely focused on translational repression during starvation, to the mechanisms governing recovery.

Our data reveal a close temporal association between metabolic changes and codon-specific ribosome pausing (**Fig. 7A, B**). Glucose repletion induced a rapid decline in AMP, coinciding with increased charging of tRNA-Ser-AGA, the only substantially undercharged serine tRNA in glucose-deprived cells, and reduced pausing at UCU and UCC serine codons. Nitrogen repletion produced a distinct response, with transiently reduced leucine tRNA charging and increased pausing at UUA and UUG leucine codons. These observations are consistent with contributions from aminoacylated tRNA availability to nutrient-specific decoding responses. Cellular nucleotide levels are known to influence translation initiation during glucose deprivation, including through the ATP-dependent activity of eIF4A^14^. Our findings suggest an additional connection between nucleotide metabolism and elongation through tRNA aminoacylation. Notably, the charging responses were not consistently explained by intracellular amino acid abundance, suggesting that additional metabolic factors influence the supply of aminoacylated tRNAs. Because elongation speed is reported to remain nearly constant across steady-state nutrient limitations^37^, these results suggest that decoding at specific codons can transiently slow during abrupt metabolic transitions, when the supply of charged tRNAs is itself changing. Codon identity may therefore be better regarded as a conditional determinant of translation whose influence depends on metabolic state.

**Fig. 7.**
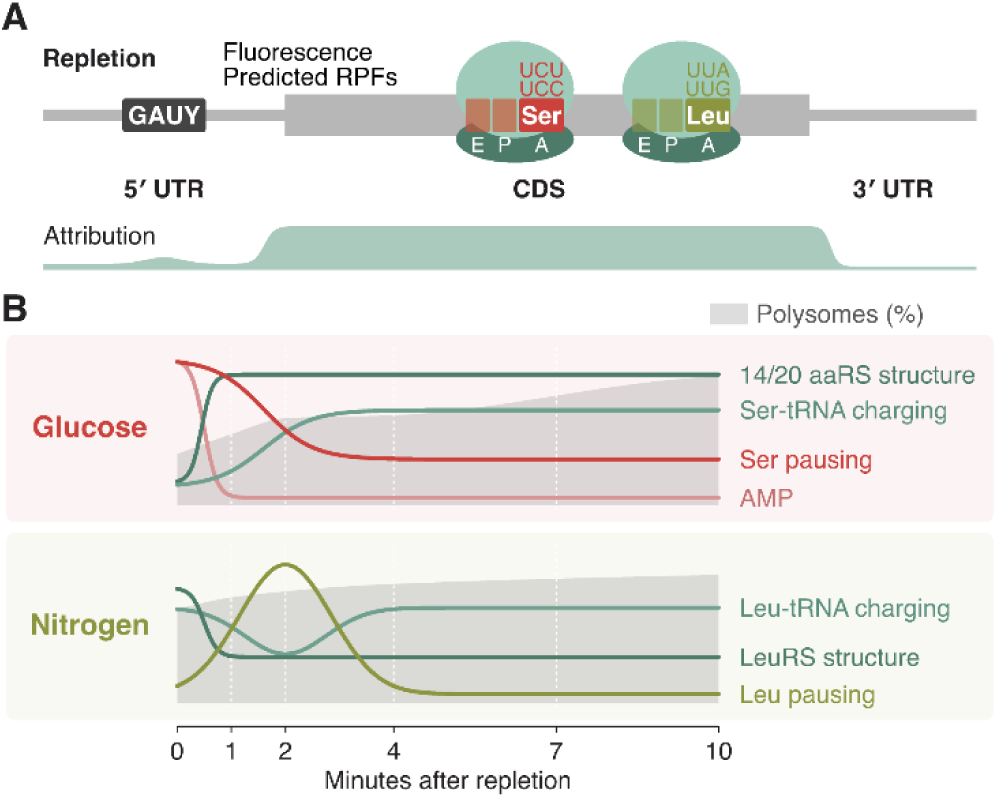
Sequence– and time-dependent translation regulation during nutrient repletion. (A) RiboNet-identified sequence features associated with ribosome occupancy. The shaded profile shows the relative contribution of transcript regions to predicted ribosome load. A-site serine and leucine codons contribute to nutrient-specific changes in ribosome pausing. (B) Schematic time courses of changes in metabolites, aminoacyl-tRNA synthetase structure, tRNA aminoacylation, and codon-specific ribosome occupancy in the first 10 minutes following glucose and nitrogen repletion. Gray shading indicates the polysome fraction.

Our structural proteomics data suggest a link between metabolism and the tRNA aminoacylation machinery. Nutrient repletion induced rapid changes in proteolytic accessibility across approximately 60% of detected translation-related proteins, including peptides from 14 of the 20 cytosolic aminoacyl-tRNA synthetases with at least one changing peptide. Several synthetase peptides correlated with cellular metabolites. Notably, a peptide in methionyl-tRNA synthetase that overlaps its ATP-binding region changed accessibility within one minute of glucose addition. Seryl-tRNA synthetase also exhibited AMP-associated structural changes during glucose repletion, while leucyl-tRNA synthetase responded to metabolites during nitrogen repletion, paralleling the codon-specific decoding responses. Aminoacyl-tRNA synthetases have previously been implicated in nutrient sensing, including the regulation of TORC1 by leucyl-tRNA synthetase^61,62^. Metabolite-associated structural states may therefore influence tRNA charging and decoding. Structural changes in eEF1A, ribosomal proteins, and other translation-associated factors indicate that metabolic adaptation extends across the translational machinery.

Translational recovery also differed in its kinetics and relative contribution of transcriptional and translational responses. Glucose repletion produced pronounced metabolic changes and rapid polysome recovery, with most changes occurring in ribosome footprints rather than mRNA abundance, accompanied by decreased pausing at a subset of serine codons. Nitrogen repletion involved substantial changes in mRNA abundance alongside translational changes, consistent with the established roles of TORC1 and the general amino acid control pathway^3,4^. Phosphate repletion altered ribosome load on many transcripts with little change in bulk polysomes or mRNA abundance, and without comparable codon-specific effects. That glucose and nitrogen repletion act predominantly at different levels indicates that the layer at which cells respond is itself nutrient-specific.

We developed RiboNet to identify sequence features associated with these nutrient-dependent translational responses. Unlike existing computational approaches, RiboNet jointly models ribosome load per transcript and codon-resolved footprint distributions, incorporating coding and untranslated mRNA regions^26–31,63,64^. A linear model of codon frequencies predicted transcript-level ribosome load as well as RiboNet, indicating that a full-sequence model matters less for predicting how many ribosomes a transcript carries than for predicting where they sit and identifying sequence features not predefined in advance. Although coding sequences contributed most strongly to predictions of ribosome load, RiboNet also identified regulatory motifs in 5′ UTRs. Among these, GAUY emerged as a conserved motif that increased reporter protein output, as demonstrated using *CPR8* and *TYW1* sequences. An independent massively parallel reporter study also identified GAUC as a sequence feature associated with translation^41^.

The increased reporter output contrasted with the reduced CDS ribosome occupancy predicted by RiboNet. One biological explanation is faster ribosome transit, which could allow greater protein production despite fewer ribosomes occupying the CDS at a given time. Differences in measurement timescales may also contribute. Ribosome profiling captures ribosome occupancy during the first 10 minutes of repletion, whereas fluorescence reflects protein accumulation over a longer period and depends on eCitrine maturation. These differences complicate direct comparisons between predicted ribosome occupancy and measured reporter output, leaving the underlying mechanism unresolved.

Our findings also raise questions about the interplay between sequence-dependent regulation and other mechanisms of translational control. Recent studies have implicated mRNA synthesis and localization in determining translational responses to acute stress^65,66^. These processes may interact with the sequence determinants identified here, whose effects depend on cellular metabolism and the availability of components of the translational machinery. Integrating sequence-based predictions with quantitative measurements offers a framework to dissect these interactions across cellular conditions. Although our study focuses on nutrient repletion in budding yeast, RiboNet is broadly applicable to other organisms and experimental conditions. Together, our findings show how changes in nutrient availability reshape translation at multiple levels, from mRNA abundance and ribosome loading to the decoding of individual codons.

## ACKNOWLEDGEMENTS

We are grateful to L. Galanti, B. Pfander, F. Bonneau and E. Conti for sharing resources, and to the Max Planck Institute of Biochemistry core facilities: NGS Core Facility for high-throughput sequencing (RRID:SCR_025746), Mass Spectrometry Facility for metabolomics and proteomics (RRID:SCR_025745), Imaging Facility for flow cytometry (RRID:SCR_025739). X.H.A. acknowledges funding from an EMBO Postdoctoral Fellowship (EMBO ALTF 792-2022) and a Research Fellowship from the Alexander von Humboldt Foundation. J.G. acknowledges funding from the European Research Council (ERC) (EPIC, Grant number: 101118521). The IT Infrastructure of the Chair of Computational Molecular Medicine was funded by the DFG under the project numbers 553375143 and 461264291. D.D.N. acknowledges funding by the Max Planck Society, the European Research Council under the European Union’s Horizon 2020 Research and Innovation Programme (ERC Starting Grant No. 803825-TransTempoFold), the EMBO Young Investigator Program (YIP 4833) and SFB 1035 (German Research Foundation DFG, Sonderforschungsbereich 1035, Projektnummer 201302640, project B14). S.M. is supported by the SciLifeLab & Wallenberg Data Driven Life Science Program (grant: KAW 2024.0159).

## AUTHOR CONTRIBUTIONS

Conceptualization: X.H.A., J.G, D.D.N; Methodology: X.H.A., S.K., D.D.N., J.G.; Software: X.H.A., F.S., B.M., S.M.; Investigation: X.H.A., K.D., B.H., S.M.; Formal analysis: X.H.A.; Resources: J.G., D.D.N.; Visualization: X.H.A.; Writing – original draft, X.H.A., D.D.N.; Writing – review & editing, S.K., S.M., J.G.; Supervision: X.H.A., J.G., D.D.N.; Funding acquisition, X.H.A., J.G., D.D.N.

## DECLARATION OF INTERESTS

D.D.N. is listed as a co-inventor on a patent application pertaining to the mim-tRNAseq technology filed by the Max Planck Society. The other authors declare no competing interests.

## DATA AVAILABILITY

All data are uploaded to public repositories and will be available upon publication.

## CODE AVAILABILITY

The RiboNet computational pipeline is available under a MIT License at https://github.com/nedialkova-lab/RiboNet. Code for reproducing all analyses and figures in this paper is accessible at https://github.com/nedialkova-lab/nutrient_repletion.

## METHODS

### Yeast strains and culture

The prototrophic S288C-derivative *S. cerevisiae* strain YSBN6^39^ (Euroscarf Y40387) was grown in synthetic defined (SD) media, containing Yeast Nitrogen Base (YNB) without amino acids (Becton Dickinson 291920) and 20 g/L glucose as a carbon source. For nutrient deprivation, the liquid media composition described in Xu et al.^40^ was used (**Supplementary Table 7**, Formedium CYN0501 and CYN6702).

For the glucose repletion, saturated cultures were diluted in SD media to an optical density 600 (OD_600_) of 0.001 and grown overnight at 30°C at 250 rpm. For nitrogen and phosphate repletion, saturated cultures were diluted in nitrogen-limiting and phosphate-limiting liquid media respectively, to an OD_600_ of 0.003 and grown overnight at 30°C at 250 rpm. This culture in limiting media before nutrient deprivation ensures the depletion of the large intracellular storage of these two nutrients^40^. Exponential phase (OD_600_ 0.5) cultures were filtered, washed with the respective nutrient-deprived media, and cultured in nutrient-deprived media for 1 hour at 30°C at 250 rpm. Cells were then harvested by rapid filtration through 0.45-µm membrane, scraping off the filter membrane and snap-freezing in liquid nitrogen, before and after 1, 2, 4, 7, 10, 14, 20, 30 minutes of adding a 5x repletion solution of glucose (final concentration 20 g/L), nitrogen (5 g/L) or phosphate (1 g/L), respectively.

For flow cytometry, YSBN6-derivative strains (**Supplementary Table 8**, see “Fluorescent reporter assay”) were pre-cultured O/N in SD media and cultured next day for two doubling times to harvest at exponential phase (OD_600_ 0.6). O/N glucose-deprivation cultures were started by diluting exponential cultures at OD_600_ 0.01. After reaching OD_600_ 0.5, cultures were harvested before and after 1 and 2 hours of adding a 5x repletion solution of glucose (final concentration 20 g/L). For harvesting, 9 mL culture was centrifuged 3000 g for 3 min and stored at –80°C for RT-qPCR, and 2 mL were collected for flow cytometry.

For immunoblotting, YSBN6-derivative strains were pre-cultured O/N in SD media and cultured next day for two doubling times to harvest at exponential phase (OD_600_ 0.6). Exactly 2 OD_600_ were harvested by centrifugation at 3000 g for 5 min.

### Calibrated ribosome profiling

#### Spike-in generation

Synthetic spike-ins were prepared as previously described^23^ with minor modifications. Two spike-in standards were generated corresponding to the *Renilla* luciferase (R_luc_) and firefly luciferase (Fl_uc_) transcripts. Synthetic mRNAs (Click Biosystems, MRNA1007 and MRNA1008) were *in-vitro*-translated using the Rabbit Reticulocyte Lysate (RRL) System (Promega L4960) to generate ribosome-mRNA complexes. Each reaction contained 50-70 μL of lysate, 2 μL of R_luc_ or 3 μL of Fluc mRNAs (500 nM), 1 μL of amino acid mixture without methionine (1mM), 1 μL of amino acid mixture without leucine (1 mM), and RNase-free water up to 100 μl. All reaction components except the mRNA were previously mixed, and the mRNA was added after denaturing at 65°C for 3 minutes. The mixture was incubated at 30°C 300 rpm for 30 minutes (Rl_uc_) or 1 hour (F_luc_), and reaction was stopped by adding an equal volume of 2x lysis buffer (for 1x: 20 mM Tris-HCl pH 8.0, 140 mM KCl, 5 mM MgCl_2_, 1% Triton X-100, 100 µg/mL cycloheximide [Sigma-Aldrich C1988], and 100 µg/mL tigecycline [Glentham Life Sciences GK6202]). The reaction mixture was directly loaded onto 10-50% sucrose gradients in 20 mM Tris-HCl pH 8.0, 150 mM KCl, 5 mM MgCl_2_, 0.5 mM DTT and 100 µg/mL cycloheximide. Gradients were centrifuged at 35,400 rpm and 4°C for 2:20 hours using a WX 90 ultracentrifuge with a TH-641 rotor (Thermo Fisher Scientific). Fractions of 290 μL were collected and those corresponding to disomes (R_luc_) or trisomes (F_luc_) were pooled. These combined fractions were aliquoted and snap-frozen using liquid nitrogen.

#### Library construction

Ribosome footprint libraries were prepared as previously described^23,24^ with minor modifications. Deprived samples at time 0 and after 1, 2, 4, 7 and 10 minutes of nutrient repletion were prepared in two biological replicates. Snap-frozen pellets were lysed in 1 mL frozen footprint lysis buffer (20 mM Tris-HCl pH 8.0, 140 mM KCl, 5 mM MgCl_2_, 1% Triton X-100, 100 µg/mL cycloheximide, and 100 µg/mL tigecycline) by cryogenic grinding using SPEX 6870 Freezer Mill or SPEX 6770 Freezer Mill (Horiba). Lysates were thawed on ice, resuspended in additional 3 mL of footprint lysis buffer, and centrifuged at 3,000 g and 4°C for 5 min. The RNA concentration of the supernatant was measured with the Qubit RNA BR kit (Invitrogen Q10210). An aliquot of 500 µg total RNA was mixed with 180µL Fluc and 160 µL Rluc spike-ins; the remaining lysate was used for polysome profiling (see “Polysome profiling by sucrose gradient”). From the spike-in-containing lysate, 10 µg RNA were used for total RNA libraries (see “Total RNA sequencing”) and the rest was filled up to 15 mL with footprint lysis buffer and layered on 3 mL sucrose cushion (1 M sucrose, 20 mM Tris-HCl pH 8.0, 150 mM KCl, 5 mM MgCl_2_, 0.5 mM DTT). Ribosomes were pelleted by centrifuging at 50,000 rpm and 4°C for 120 minutes, and carefully washed and resuspended in 500 µL footprint lysis buffer without antibiotics by intermittent pipetting on ice. The RNA concentration was measured with the Qubit RNA HS kit (Invitrogen Q32852) and 350 µg RNA in 400 µL resuspension volume was digested with 1,000 U Ambion RNase I (Invitrogen, AM2295) shaking at 500 rpm and 25°C for 1 hour. If the most diluted sample of the batch had lower than 350 µg in 400 µL, the RNA amount present in 400 µL of this sample was used and all other samples diluted to the same concentration. Digestion was stopped with 200 U SUPERase·In (Invitrogen AM2696) and directly loaded on a 10-50% sucrose gradient in 20 mM Tris-HCl pH 8.0, 150 mM KCl, 5 mM MgCl2, 0.5 mM DTT. Gradients were centrifuged at 38,000 rpm and 4°C for 1:40 hours using a WX 90 ultracentrifuge with a TH-641 rotor (Thermo Fisher Scientific). Fractions of 290 μL were collected and those corresponding to monosomes were pooled, mixed with 1% SDS and snap-frozen in liquid nitrogen.

For RNA isolation, samples were thawed for 1 minute in 65°C waterbath then mixed with 1 volume of 65°C acid phenol (pH 4.3) and vortexed vigorously followed by incubation at 65°C for 5 min with intermittent mixing. After addition of 1/9 volume 1-bromo-3-chloropropane (BCP, Sigma Aldrich), samples were centrifuged at 10,000 g for 5 min and the aqueous phase was transferred to a new tube. After an additional round of hot acid phenol/BCP and a round of BCP-only extraction, RNA was precipitated from the aqueous phase by the addition of 2.5 volumes of ice-cold 100% ethanol, 1/10 volume of 3 M NaOAc, and 50 µg glycogen (Ambion AM9510). After precipitation (for at least 30 min at –20°C), samples were centrifuged at 16,000 g and 4°C for 30 minutes, pellets were briefly air-dried and resuspended in RNase-free water. 5 µg extracted RNA was resolved on a denaturing 15% polyacrylamide/7M urea/1xTBE gel alongside 19-, 22-, 26-, and 32-nt RNA markers and visualized with SYBR Gold, excising the region corresponding to 19-32 nt. Gel slices were crushed with disposable pestles, soaked in 400 µL of gel elution buffer (0.3 M NaOAc pH 4.5, 0.25% SDS, 1 mM EDTA pH 8.0), incubated at 65°C for 10 min, snap-frozen on dry ice, and thawed at 65°C for 5 min. RNA was eluted overnight at room temperature with continuous mixing. Gel pieces were removed with Costar Spin-X centrifuge tube filters and RNA was precipitated from the flow-through with 2.5 volumes 100% ethanol and 35 μg of glycogen. RNA 3′ ends were dephosphorylated with T4 PNK (New England Biolabs M0201L) for 45 min at 37°C and ethanol-precipitated in the presence of 10 μl of 3M NaOAc (pH 4.5) and 35 μg glycogen. Pre-adenylated adapters with 5 randomized nucleotides at their 5′ end were then 3′-ligated with T4 ligase 2, truncated KQ (New England Biolabs M0373S) at 25°C for 3 hours. Six ligation reactions of adapters with different indices were pooled and purified (Oligo Clean & Concentrator kit, Zymo Research D4060). Eluted RNA was resolved on a denaturing 12% polyacrylamide/7M urea/1xTBE gel alongside Low Range ssRNA marker (New England Biolabs) and visualized with SYBR Gold. The linker-ligated fragments of ∼50-60 nt were excised, crushed and eluted from the gel. cDNA was synthesized by reverse transcription by hybridizing RNAs at 65°C for 5 min with a DNA oligo complementary to the 3′ 15 nt of the adapter sequence and with two 5′-RN to minimize circularization biases, cooling down on ice, and incubating at 50°C for 30 min with 0.5 mM dNTPs, 10 mM DTT, 20 U SUPERase·In, and 1x Protoscript buffer and 50 U ProtoScript II (New England Biolabs M0368L). The reaction was resolved on a denaturing 10% polyacrylamide/7M urea/1xTBE gel, visualized with SYBR Gold, and the highest band at ∼100-110 nt was excised, crushed, eluted, and ethanol-precipitated in the presence of 50 µg glycogen and 30 µL 3M NaOAc (pH 4.5). cDNA was circularized with TS2126 RNA ligase 1^67,68^ (a.k.a. CircLigase) in circularization buffer (10 mM KCl, 5 mM MgCl_2_, 2.5 mM MnCl_2_, 1 M Betaine and 50 mM MOPS pH 7.5, 50 µM ATP, 1 mM DTT) at 60°C for 3 hours followed by 10 min at 80°C to stop the reaction. Circularized cDNA was rRNA-depleted by hybridization of 0.5 μM biotinylated oligos complementary to the 12 most common rRNA contaminants in our workflow, followed by two rounds of incubation and magnetic beads removal with Dynabeads MyOne Streptavidin C1 (0.5 mg beads per round, Invitrogen 65001). Eluate was purified (Oligo Clean & Concentrator kit) and 1/4 of sample amplified by library PCR (KAPA HiFi Hotstart PCR Kit, Roche 7958897001) with 8 cycles. Reactions were resolved on a 8% polyacrylamide/1xTBE gel, visualized with SYBR Gold, and dsDNA of ∼160 nt was excised, crushed, eluted, and ethanol-precipitated in the presence of 50 µg glycogen and 40 µL 3M NaOAc (pH 4.5). Libraries were quantified with the Qubit dsDNA HS kit (Invitrogen Q32854) and 120-bp single-end sequencing was performed on a NovaSeq 6000 platform (Illumina 20028401).

#### Data analysis

The reference genome annotation of *S. cerevisiae* strain S228C was downloaded from the Saccharomyces Genome Database^69^ (SGD, genome assembly R64-2-1). The annotated GFF features were filtered to remove “Dubious”-labeled ORFs, resulting in a total of 5,913 annotated genes. This genome assembly and annotation was merged with the transcript sequences and annotations of synthetic spike-ins Fluc and Rluc. This combined reference genome was used for ribosome profiling and matched RNA-seq read alignment. Genome index was generated using STAR^70^ (v2.7.11a) with the arguments “—genomeSAindexNbases 10 –-sjdbGTFfeatureExon CDS –-sjdbGTFtagExonParentTranscript Parent –-sjdbOverhang 35”.

Sequencing libraries were demultiplexed and adaptor-trimmed with Cutadapt^71^ (v4.9) with the following arguments: “--no-indels –q 30,30 –-trimmed-only –m 10”. Reads were further trimmed to remove low quality bases, the five linker-ligated 3′-NNNNN, and the two 5′-RN nucleotides introduced in RT with “-j 40 –m 10 –u 2 –u –5”. Trimmed reads were aligned to the S288C non-coding RNA reference from SGD^69^ using bowtie^72^ (v1.3.1) with “-p 40 –S –-best”, and ncRNA-unmapped reads were aligned to the reference genome above using STAR^70^ with the arguments: “--outFilterMultimapNmax 1 –-outSAMtype BAM SortedByCoordinate –– outFilterMismatchNmax 0 –-alignEndsType Local –-seedSearchStartLmax 14 –-alignIntronMax 10000 –-outFilterIntronMotifs RemoveNoncanonicalUnannotated –-quantMode TranscriptomeSAM –-limitBAMsortRAM 5000000000 –-outSAMattributes NH HI AS nM NM MD”.

The QC of ribosome profiling libraries, including footprint length distribution and 3-nt periodicity, was performed using the preprocessing module of the developed RiboNet package. A-site offset determination is performed using the Integer Programming method implemented in RiboA^73,74^ (v1.0.2). Read lengths corresponding to ribosomes with vacant (20-23 nt) or occupied (27-32 nt) ribosomal A sites were processed separately, and total read counts per gene aggregated for differential expression analysis. Differentially expressed genes along the 10-min time course were determined with *limma-voom*^75^ (v3.58.1) using a cubic spline (knots at 0.5, 2.5 and 5 min), and applying normalization for negative control samples with RUVseq^76^ (v1.36.0).

#### Calibration with spike-ins

For calibration of the absolute number of ribosomes per transcript using spike-ins, we used the calculation explained in Tomuro et al^23^. Briefly, ribosomes per transcript were computed as the ratio of ribosome footprint abundance (RPM) to mRNA abundance (TPM) for each gene. These ratios were then calibrated to absolute values using the spike-in reporters of known ribosome occupancy, Rluc disomes (2 ribosomes per transcript) and Fluc trisomes (3 ribosomes per transcript). For each sample, a scaling factor was set so that the summed Rluc and Fluc ratios equal 5 (i.e., 5 / (RPF_Fluc_/RNA_Fluc_ + RPF_Rluc_/RNA_Rluc_)), and all genes in that sample were multiplied by it. The amount of spike-in reads per sample was 0.03% on average. Significant changes in log-transformed ribosomes per transcript were determined with *limma*^75^ using a cubic spline (knots at 0.5, 2.5 and 5 min). Significant genes were clustered by time pattern using 1 − *R_Pearson_* as distance metric and Ward’s minimum variance linkage method. For each cluster, GO enrichment analysis of biological processes was performed with clusterProfiler^77^ (v4.10.1).

#### Codon occupancy and elongation time estimates

The ribosome occupancies on each of the 61 amino-acid-encoding codons were computed as the ratio between the ribosome density within each A-site codon and the overall density along each gene CDS (excluding the first and last 15 nucleotides), and taking the average across all genes. This normalized occupancy reflects the relative decoding time of translation elongation at each codon^29^. The geometric mean of all codons of a gene therefore corresponds to an estimate of ribosome elongation time^42^.

### Total RNA sequencing

#### Library construction

For normalizing ribosome profiling libraries to total RNA abundance, one replicate of each deprived sample at time 0 and after 1, 2, 4, 7 and 10 minutes of nutrient repletion was analyzed. An aliquot of 10 µg RNA of the same yeast cell lysate used for ribosome profiling, which contains the spike-ins, was resuspended in 600 µL RNA isolation buffer (50 mM NaOAc pH 4.5, 10 mM EDTA pH 8.0, 1% SDS). An equal volume of hot acid phenol (pH 4.3) was added, and the cell suspension was vortexed vigorously followed by incubation at 65°C for 5 min with intermittent mixing. After addition of 75 µL BCP (Sigma Aldrich), samples were centrifuged at 10,000 g for 5 min and the aqueous phase was transferred to a new tube. After an additional round of hot acid phenol/BCP and a round of BCP-only extraction, RNA was precipitated from the aqueous phase by the addition of 2.5 volumes of ice-cold 100% ethanol and 25 µg glycogen (Ambion). After precipitation (for at least 30 min at –20°C), samples were centrifuged at 16,000 g and 4°C for 30 minutes, pellets were briefly air-dried and resuspended in RNase-free water. The extracted RNA was DNAse-treated and purified (RNA Clean & Concentrator-5, Zymo Research R1013). rRNA was depleted by hybridization of *S. cerevisiae* riboPOOLs (siTOOLs Biotech dp-P012-5), followed by two rounds of incubation and magnetic beads removal with Hydrophilic Streptavidin Magnetic Beads (0.5 mg beads per round, New England Biolabs S1421S). The final supernatant was ethanol-precipitated and resuspended in RNAse-free water. 300 ng of rRNA-depleted RNA was used for library preparation with the Zymo-Seq RiboFree Total RNA Library Kit (Zymo Research R3003), skipping the universal depletion steps as indicated in the manufacturer’s protocol appendix D and with a final library PCR of 13 cycles. The fragment size of final libraries was determined on an Agilent TapeStation. Libraries were quantified with the Qubit dsDNA HS kit and 120-bp single-end sequencing was performed on a NovaSeq 6000 platform.

#### Data analysis

The genome index for RNA-seq alignment was generated as described for ribosome profiling, with the STAR^70^ arguments “--genomeSAindexNbases 10 –-sjdbGTFfeatureExon CDS –-sjdbGTFtagExonParentTranscript Parent –-sjdbOverhang 80”. Potential 3′ adaptors in RNA-seq reads were removed using TrimGalore^78^ (v0.6.4) with default settings. Trimmed reads were processed as described with ribosome profiling libraries, filtering ncRNA-mapping reads with bowtie^72^ and aligning the remaining reads with STAR^70^ (“--outFilterMultimapNmax 1 –-outSAMtype BAM SortedByCoordinate –-outFilterMismatchNmax 1 –-quantMode TranscriptomeSAM –-limitBAMsortRAM 50000000000”). Read counts per gene were then generated using featureCounts^79^ (v2.0.6), with arguments “-B –g Parent –t CDS”. Differentially expressed genes along the 10-min time course were determined with *limma-voom*^75^ using a cubic spline (knots at 0.5, 2.5 and 5 min).

### Deep learning modeling of mRNA translation

#### Data preprocessing

The input of RiboNet is a sample file identifying all BAM alignment files and metadata of ribosome profiling samples, the RNA-seq counts per gene, and the reference genome FASTA and annotation GFF (see previous sections). The sample file defined selected footprint lengths of 20-23 nucleotides for short footprints and 27-32 nucleotides for long footprints. Annotated 5′ UTRs and 3′ UTRs can be provided explicitly if available; the S288C UTR annotations were downloaded from SGD^69^ (http://sgd-archive.yeastgenome.org/sequence/S288C_reference/). The “ribonet preprocessing” command was executed with the following arguments: –-spikein Fluc,Rluc –-spikein_absolute_ribosomes 5 –-utr5annot SGD_all_ORFs_5prime_UTRs.fsa –-utr3annot SGD_all_ORFs_3prime_UTRs.fsa

This first step of RiboNet generates a QC report for ribosome profiling data, determines the A-site offset using RiboA^73,74^, computes spike-in calibration, and extracts all transcript sequences and ribosomal footprints distribution. Applying the default RiboNet filters of sequence length and minimum average footprint density of 0.1 reads per codon, data resulted in a total of 4799, 4651, and 4677 genes for glucose, nitrogen and phosphate repletion datasets, respectively.

#### RiboNet architecture and training

RiboNet is a sequence-to-function model, which predicts the ribosome counts and distribution along the transcript from sequence (**Supplementary Table 9**). Although different architectures are available in the developed package (see package documentation), we used here a model that predicts the distribution of ribosome footprints at codon resolution. The “ribonet train” command was executed with the following arguments:

--lm5 specieslm5 –-species_proxy5 kazachstania_africana_cbs_2517_gca_000304475 –-lm3 specieslm3 –-species_proxy3 candida_glabrata

In this configuration, 5′ UTR and 3′ UTR sequences are tokenized and processed through the DNA language model SpeciesLM^44^ (pre-trained with 1,500 fungal genomes, using as proxy the closest species to *S. cerevisiae* in the training set). A weighted average of all layers of the frozen embeddings is retrieved (weight parameters fitted during training), average-pooled to 30% of the original length and projected down to 128 dimensions. The coding sequence (CDS) is one-hot encoded at codon resolution (non-overlapping 3-mers), and processed through a 1D-convolutional block of 5 layers. The resulting 128-dimensional 5′ UTR, CDS, and 3′ UTR embeddings are then concatenated, added with a positional embedding, and processed through one layer of transformer encoder to capture long-range effects. These embeddings are used to generate two separate heads in a multi-task manner (i.e. each task corresponding to each condition), similar to the BPNet architecture^80^. The first head consists of a 1D-convolutional layer that predicts the distribution of ribosome footprints along the transcript. The second head performs average-pooling of the embeddings and predicts the ribosomes per transcript through a linear layer, which are then normalized to the RNA abundance to output the total ribosome footprint counts per gene. To train this model, we defined a BPNet-like loss function which combines both distribution and counts predictions.

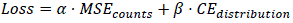

Where *α* and *β* were set to 0.1 and 1 (defaults in RiboNet), which weight the contribution of the Mean Squared Error of counts and the Cross Entropy of distribution probabilities, respectively. The loss function is computed for each biological replicate independently and averaged, so that the model can assess the variability between replicates. Genes in the original datasets were split into 70% training set, 15% validation set, and 15% test set. The model was trained with the default parameters: batch size of 32, learning rate of 0.0005, multi-step scheduler with 5-epoch warm-up, maximum of 50 epochs and patience of 10 epochs.

This training step generated a training report containing the following evaluation metrics of the model on the test set: percentage of total counts variance explained, correlation of predicted versus observed counts per condition, correlation of predicted versus observed footprint distribution per position, and normalized codon occupancies in predicted and observed footprint distributions.

#### Model interpretation

Identification of regulatory features of RiboNet was performed by *in silico* mutagenesis (ISM). For efficient parallelisation, this step is divided into three steps in the RiboNet package. First, “ribonet interpret” initializes the scoring parameters: “--save-distribution”, which enables the saving of both counts and distribution, and “--n_batches”, which splits ISM in several batches of genes for parallelisation. For each batch, the command “score batch –-batch N” mutates every position of each transcript to all other nucleotide possibilities and performs a forward pass through the model to measure the change in counts and distribution of each mutation. Finally, “ribonet report” processes the ISM results to identify regulatory motifs, which was executed with the argument “--control” set to the corresponding starting time point of each nutrient repletion. For each nucleotide position, absolute attribution is scored as *max*(|*ln WT* − *ln Mut*|) and relative attribution compared to the specified control is computed as *avg*[(*ln WT* − *ln Mut*)*_cond_* − (*ln WT* − *ln Mut*)*_control_*].

Interpretation analysis of the counts head consists of TF-MoDISco^50^ (v2.2.1) analysis to identify recurrent sequence motifs (“seqlets”) with high attribution score, both in absolute terms and in comparison to the control condition. For footprint distribution effects, RiboNet identifies differentially occupied P-sites between conditions and reports the attribution scores of a nucleotide window around them. All these results are saved and printed in an interpretation report. If both long and short footprints are defined in the sample file, the interpretation analyses are also performed independently for each footprint length.

### Benchmarking of RiboNet models

To benchmark the contribution of individual input modalities to ribosome profiling prediction, we trained five model variants of RiboNet across three nutrient-limitation datasets (glucose, nitrogen and phosphate), which included: (i) full RiboNet containing 5′ UTR, CDS and 3′ UTR; (ii) 5′ UTR ablation, containing only CDS and 3′ UTR; (iii) 3′ UTR ablation, containing only 5′ UTR and CDS; (iv) both UTRs ablation, containing only CDS; (v) counts-only, as the full model but trained with the profile-loss weight β = 0. For each dataset and variant, 50 independent training runs were performed, each using different train/validation/test gene sets (70%/15%/15% split). In addition to the RiboNet models, two linear baselines were fitted on the training set and evaluated on the test set: (i) RNA baseline as an univariate linear regression of length-normalised log RPM on log(RNA-seq TPM+1); (ii) codon usage baseline as a no-intercept linear regression of RNA– and length-normalized log RPM on the 61 codon frequencies without intercept.

Model performance was computed over all replicates and all genes of the held-out test set for short (20–23 nt) and long (27–32 nt) RPF read-length classes separately. Three metrics were quantified: (a) fraction of variance in log RPM explained (EV%), (b) Pearson correlation coefficient across genes and taking the median of all time points, (c) Pearson correlation coefficient across time points and taking the median of all genes.

### Evolutionary conservation of 5′ UTR motifs

We first identified the gene families for all genes across 1,500 fungal genomes using the homology inference module of FastOMA^81^ (v0.5.1). This process began with the rapid k-mer-based assignment of genes to known OMA gene families using OMAmer^82^ (v2.1.2), followed by *de novo* inference of gene families for the remaining genes using Diamond DeepClust^83^ (v2.1). The resulting assignments constitute the set of homology families across the 1,500 fungal genomes.

For each GAUY seqlet identified by TF-MoDISco^50^, we extracted all high-confidence seqlet-containing genes to compute the evolutionary conservation of the corresponding motif. For each gene, the 200 nucleotides upstream of the start codon of all fungi homologs were retrieved and motif enrichment was computed using Simple Enrichment Analysis (SEA) from the MEME suite^59^ (v5.5.9). In homology families larger than 2,000 genes, a random subset of 2,000 homologs was used. Motif enrichment was computed among species sets at different evolutionary distances within the fungi phylogenetic tree from NCBI. The NCBI tree was acquired using the NCBITaxa submodule of the ETE3 package^84,85^. In parallel for each gene and evolutionary distance set, background 5′ UTR conservation was computed as the median enrichment of 100 random 5′ UTR k-mers of the same size (filtered for overlapping sequences and duplicated sequences).

To infer orthology relationships, we ran FastOMA^81^ with default parameters on 1,500 fungal genomes and extracted all genes that are orthologous to CPR8 and TYW1. The same SEA was performed among orthologs of CPR8 and TYW1, measuring the enrichment of the GAUUAU and GAUCA motifs, respectively.

### Structural analysis of 5′ UTR motifs

To test whether sequence flanking the discovered GAUY motif tends to fold into an RNA stem-loop, we performed two complementary analyses. For each of the high-confidence GAUY-containing genes, the 10 nt immediately upstream and downstream of the GAUY motif were extracted. Stem potential was scored as the longest run of stacked antiparallel base pairs between the upstream flank and the reversed downstream flank, allowing Watson–Crick and G·U wobble pairs. As a matched control, we sampled, per gene, 50 random same-shaped windows (10 + motif length + 10 nt) from that gene’s own 5′ UTR sequence and computed the same longest-stem statistic.

For the targeted structural analysis of CPR8 and TYW1 5′ UTRs, we retrieved 100 nt upstream of the CDS start for each gene’s orthologs, restricted them to the ∼145-Mya clade, and aligned them with LocARNA^60^ “mlocarna” (v2.0.1) with default parameters and specifying the NCBI fungi phylogenetic tree.

### Proteomics

#### Sample preparation

Yeast cells exponentially growing in SD media and after one hour of each nutrient deprivation (corresponding to time 0 of the time course) were prepared in triplicates. Snap-frozen pellets were resuspended in 1 mL of preheated SDC buffer containing 1% sodium deoxycholate (SDC, Sigma-Aldrich), 40 mM 2-chloroacetamide (CAA, Sigma-Aldrich), 10 mM tris(2-carboxyethyl)phosphine (TCEP, Thermo Fisher Scientific), and 100 mM Tris (pH 8.0). Samples were incubated at 95°C for 5 minutes, followed by ultrasonication in an ice bath. The heating and ultrasonication steps were repeated once. Next, 10 µL of lysate from each sample was transferred to a 96-well plate for automated processing using the APP96 robotic platform (PreOmics P.O.00139) and the iST APP96 96x sample kit (PreOmics P.O.00141). The automated workflow included the addition of 200 µL LYSE buffer. After incubation at 80°C for 10 minutes with shaking, the plate was cooled to 37°C and DIGEST buffer containing Lys-C and trypsin was added. Samples were incubated at 37°C for 1 hour, and digestion was terminated by adding 100 µL STOP buffer. Following thorough mixing, samples were loaded onto PreOmics cleanup cartridges (POPTips), which were washed sequentially with 200 µL of Wash Buffers I and II. Peptides were eluted with 165 µL ELUTE buffer, vacuum-dried, and 200 ng of peptide material was loaded onto Evotips (Evotip Pure, Evosep).

#### LC-MS/MS data acquisition

Peptides were eluted from Evotips onto a 15 cm PepSep C18 column (1.5 µm; Bruker Daltonics) using an Evosep One HPLC system (Evosep). The column temperature was maintained at 50°C, and peptide separation was performed using the 30 SPD method. Eluted peptides were directly ionized and introduced into a timsTOF Pro mass spectrometer (Bruker) via electrospray ionization. Data was acquired in data-independent acquisition (DIA)-PASEF mode using timsControl software. Mass spectra were acquired over an m/z range of 100-1700 with an ion mobility range of 1/K_0_ = 0.70-1.30 Vs·cm^-2^. Equal ion accumulation and ramp times of 100 ms were applied in the dual TIMS analyzer, resulting in a spectral acquisition rate of 9.52 Hz. DIA-PASEF scans covered a mass range of 350.2-1199.9 Da with an ion mobility range of 1/K_0_ = 0.70-1.30 Vs·cm^-2^. Collision energy was linearly ramped according to ion mobility, from 45 eV at 1/K_0_ = 1.30 Vs·cm^-2^ to 27 eV at 1/K_0_ = 0.85 Vs·cm^-2^. A total of 42 DIA-PASEF windows were acquired per TIMS scan with alternating precursor isolation windows, resulting in an estimated cycle time of 2.21 seconds.

#### Data analysis

Raw data was processed using Spectronaut (Biognosys, v18.0.230605.50606) in directDIA+ (library-free) mode with default settings. Spectra were searched against a predicted yeast protein library derived from the UniProt database (Swiss-Prot and TrEMBL; downloaded in 2023). Carbamidomethylation of cysteine residues was specified as a fixed modification, while methionine oxidation and protein N-terminal acetylation were included as variable modifications. Protein quantification across samples was performed using label-free quantification (MaxLFQ) at the MS^2^ level.

Significant changes in log-transformed protein abundances between nutrient deprivation conditions were determined with *limma*^75^. GO enrichment analysis of biological processes was performed with clusterProfiler^77^.

### Limited proteolysis coupled to mass spectrometry

#### Sample preparation

LiP-MS samples were prepared as previously described^49^. Deprived samples at time 0 and after 1, 2, 4, 7, 10, 14, and 20 minutes of nutrient repletion were prepared in triplicates. Snap-frozen pellets were lysed in 250 µL LiP buffer (20 mM HEPES-KOH pH 7.5, 150 mM KCl, 10 mM MgCl2, 1x cOmplete EDTA-free protease inhibitor cocktail [Roche 11873580001]) by cryogenic grinding using SPEX 6870 Freezer Mill (Horiba). Lysates were kept frozen at – 80°C. All time points and replicates were randomized to minimize batch effects and processed further in 4 separate days. Powdered lysates were thawed on ice, briefly centrifuged, transferred to pre-chilled 1.5 mL non-stick tubes, and clarified twice by centrifugation at 16,000 g for 5 min at 4°C, transferring the supernatant to a fresh pre-chilled tube after each spin. Protein content of extracts was quantified by BCA assay (Thermo Fisher Scientific 23225), adjusted to 2 mg/mL in LiP buffer, and aliquoted in 100 µg into non-stick PCR strip tubes for limited proteolysis (LiP) and for trypsin-only control (TrP). PCR strips were processed in a Thermocycler block, starting with 5 min equilibration at 25°C followed with either 5 µL water (TrP) or 5 µL PK solution (LiP, 0.2 µg/µL in LiP buffer) was added (enzyme:substrate 1:100, w/w) and incubated for exactly 5 min at 25°C. Reactions were stopped by heating at 99°C for 5 min, cooling down to 4°C, and adding 55 µL DOC (10% w/v, freshly prepared). Samples were stored at –20°C.

LiP samples were further processed using the PreOmics APP96 automation platform (PreOmics). Lysates (10 μl) were loaded into a 96-well plate, followed by addition of 200 μL LYSE buffer. Samples were incubated at 80 °C for 10 min with shaking, cooled to 37 °C, and digested with DIGEST buffer containing LysC and trypsin for 1 h at 37 °C. Digestion was stopped by addition of 100 μl STOP buffer. Following mixing by pipetting, samples were loaded onto PreOmics POPTIPS, washed sequentially with 200 μl Wash Buffer I and II, and eluted in 165 μl ELUTE buffer. Eluates were vacuum-dried and reconstituted in 50 μl LC-load buffer before loading onto Evotip Pure tips (Evosep).

TrP samples were heated at 95 °C for 5 min and sonicated using a Bioruptor Plus system (Diagenode) for 10 cycles of 30 s at high intensity. Heating and sonication were then repeated. Samples were diluted 1:1 with MS-grade water (VWR) and digested overnight at 37 °C with 1 μg Lys-C (Wako) and 1.5 μg trypsin (Promega). Digestion was stopped by acidification to 1% trifluoroacetic acid (TFA; Merck), and peptides were loaded onto Evotip Pure tips (Evosep).

For spectral library generation, equal volumes of LiP samples were pooled and fractionated into 32 fractions by high-pH reversed-phase chromatography on a Vanquish Neo system using a BEH C18 column (300 Å, 1.7 μm, 300 μm × 150 mm; Waters). Peptides were separated using a gradient from buffer A (20 mM ammonium bicarbonate in water, pH 8.8) to buffer B (20 mM ammonium bicarbonate in 80% acetonitrile, pH 8.8). Fractions were vacuum-dried, reconstituted in 0.1 % formic acid and loaded on Evotip Pure tips (Evosep).

#### LC-MS/MS data acquisition

Evotips were analyzed on an Evosep One HPLC system using a C18 column (PepSep C18, 15 cm × 150 μm, 1.5 μm; Bruker Daltonics) maintained at 50 °C and operated using the 30 samples-per-day (SPD) method. Eluting peptides were directly introduced into an Orbitrap Exploris 480 mass spectrometer (Thermo Fisher Scientific).

For DIA, full MS scans were acquired from 300–1650 m/z at a resolution of 120,000, with an AGC target of 300% and a maximum injection time of 20 ms. Precursors were fragmented by HCD at normalized collision energies of 25, 27.5, and 30%, and fragment ions were acquired in 44 DIA windows at a resolution of 30,000 with an AGC target of 300%.

For DDA, used for spectral library generation, full MS scans were acquired from 300–1650 m/z at a resolution of 60,000. The 15 most intense precursors were selected for HCD fragmentation at a normalized collision energy of 28. MS2 spectra were acquired at a resolution of 15,000. AGC targets were 300% and 100% for MS1 and MS2 scans, respectively, with maximum injection times of 25 and 28 ms. Dynamic exclusion was set to 30 ms.

#### Data analysis

Raw data were processed using Spectronaut 18.0.230605.50606 (Biognosys). A spectral library for the LiP-MS samples was generated from the high-pH fractionated samples acquired in DDA mode using Pulsar with default settings, except that searches were performed against the *S. cerevisiae* UniProt database (downloaded in 2023), with Trypsin/P specified as the enzyme and semi-specific digestion enabled. A hybrid library was then generated in Pulsar by combining the DIA runs analyzed using directDIA+ with the DDA-derived spectral library described above. This hybrid library was used for all library-based analysis of both LiP and TrP samples with default settings, except that semi-specific digestion was specified for LiP samples and the minimum peptide length was set to six amino acids.

Results were exported using the LiPAnalyzeR Spectronaut scheme format^86^, and “PEP.Quantity” and “PG.Quantity” were extracted to measure peptide and protein quantities, respectively. Measurements were log_2_-transformed and filtered for values with a log_2_-abundance above 5. Batch-correction was applied using the “removeBatchEffect” function of *limma*^75^, which considers both the biological design and the batch effect related to different PCR strips during sample preparation.

For protein abundance, only proteins with at least one time point with 2/3 measurements and at least 4 measurements in total per nutrient repletion set were kept. This resulted in a high-confidence dataset of 2,665 proteins in glucose repletion, 2,676 proteins in nitrogen repletion, and 2,668 proteins in phosphate repletion. Significant changes in protein abundance along the 20-min time course were determined with *limma*^75^ using a cubic spline (knots at 0.5, 2.5 and 5 min).

Structural changes of limited-proteolysis peptides were analyzed using LiPAnalyzer^86^, removing unwanted variance of both tryptic and half-tryptic peptides with a “LiPPep ∼ TrPProt” model (“FTHTjoin” mode). The residuals of this model were filtered similarly to protein abundance, keeping only peptides with at least one time point with 2/3 measurements and at least 4 measurements in total, resulting in a high-confidence dataset of 25,723 peptides (2,457 proteins) in glucose repletion, 27,180 peptides (2,496 proteins) in nitrogen repletion, and 26,463 peptides (2,479 proteins) in phosphate repletion. Filtered residuals were analyzed with *limma*^75^ using a cubic spline (knots at 0.5, 2.5 and 5 min). Significant peptides were clustered by time pattern using 1 − *abs*(*R_Pearson_*) as distance metric and Ward’s minimum variance linkage method.

#### Peptide mapping to three-dimensional protein structures

Three-dimensional protein structures were obtained from the RCSB Protein Data Bank or the AlphaFold EBI database. Residue-level mapping between UniProt sequence positions and PDB chain/residue identifiers was performed using the PDBe SIFTS API. Structures were rendered using py3Dmol: protein chains are displayed as Van der Waals surfaces and nucleic acid chains as cartoons. Residues covered by significant LiP-MS peptides (adj. p < 0.05 in at least one condition) are highlighted, and those also significantly correlated with metabolite levels shown in a darker color.

### Metabolomics

#### Sample preparation

Deprived samples at time 0 and after 1, 2, 4, 7, 10, 14, 20, and 30 minutes of nutrient repletion were prepared in triplicates. Samples for metabolomics were collected using the cold methanol quenching-boiling ethanol extraction method (CMBE)^87^. 10 mL of yeast cell culture were quenched in 20 mL of cold methanol, kept on dry ice, and subsequently centrifuged at 3,220 g and –9°C for 3 min to discard the supernatant. The pellet was resuspended and vortexed in 1.5 mL of preheated (75°C) ethanol solution (75% v/v) and 50 μL 25 μM ^13^C^15^N-labeled amino acid mix as internal standards (Cambridge isotopes MSK-A2-1.2). Metabolites were extracted at 95°C for 3 min in a water bath followed by 10 min chilling on ice. After centrifuging at 4,000 rpm and 4°C for 5 min, the supernatant was collected in a 2-mL tube, dried in a vacuum concentrator at 30°C, and stored at –80°C.

#### LC-MS/MS data acquisition

Data acquisition was performed using a Vanquish Flex UHPLC system (Thermo Fisher Scientific) coupled to a Q Exactive HF mass spectrometer (Thermo Fisher Scientific). Vacuum-dried metabolite extracts were reconstituted in 100 µL of buffer A (0.1% formic acid), and 3 µL of each sample was injected onto a Kinetex F5 column (2.1 × 100 mm, 2.6 µm; Phenomenex) operated at a flow rate of 200 µL/min. Chromatographic separation was achieved using buffer B (100% acetonitrile, 0.1% formic acid), which was held at 0% for 2 minutes, increased to 95% over 12 minutes, and maintained at 95% for an additional 2 minutes. The column was subsequently re-equilibrated at 0% buffer B for 4 minutes. The mass spectrometer was operated in positive ionization mode with data-dependent acquisition. Full MS^1^ scans were acquired over an m/z range of 75-1000 at a resolution of 120,000. Heated electrospray ionization (HESI) source parameters were as follows: sheath gas (N_2_) flow rate of 47, auxiliary gas flow rate of 10, and sweep gas flow rate of 2 (all in arbitrary units). The spray voltage was set to 3.20 kV, the capillary temperature to 250°C, and the auxiliary gas heater temperature to 380°C. The five most intense precursor ions were selected for fragmentation by higher-energy collisional dissociation (stepped HCD with normalized collision energies of 20, 40, and 60). MS^2^ spectra were acquired at a resolution of 30,000. The automatic gain control (AGC) targets were set to 3 × 10^6^ for MS^1^ and 1 × 10^5^ for MS^2^, with maximum injection times of 200 ms and 50 ms, respectively.

#### Data analysis

Untargeted metabolomics data were processed using MASSter^88^ (Zamboni Lab, ETH Zürich, v0.11.17), with the default parameters for 20-minute gradient, Orbitrap and positive acquisition mode. This workflow included feature detection, consensus feature alignment across samples, and reproducible quantification. Compound identification was performed with tima^89^ (v2.12.0.9000), imported into the MASSter workflow, and combined with the MS1-level compound identification of MASSter for the remaining unknown features. For each nutrient repletion set, only features with at least one time point with 2/3 measurements and at least 4 measurements in total were kept, which resulted in a high-confidence dataset of 1,859 features in glucose repletion (493 MS^1^– and 72 MS^2^-identified compounds), 1,838 features in nitrogen repletion (489 MS^1^– and 71 MS^2^-identified compounds), and 1,841 features in phosphate repletion (488 MS^1^– and 71 MS^2^-identified compounds). Quantification measurements were normalized to the median log_2_-abundance of internal standards followed by probabilistic quotient normalization (PQN). Time-course replicates were batch corrected using the “removeBatchEffect” function of *limma*^75^, considering also the biological design.

Significant changes in log_2_-transformed metabolite abundances along the 30-min time course were determined with *limma*^75^ using a cubic spline (knots at 0.5, 2.5 and 5 min). Significant metabolites were clustered by time pattern using 1 − *R_Pearson_* as distance metric and Ward’s minimum variance linkage method. For each cluster, peak set enrichment analysis of KEGG metabolic pathways was performed with the mummichog algorithm of the MetaboAnalystR^90^ package (v4.3.0).

For targeted data analysis, peak areas were identified and extracted based on accurate MS^1^ masses and retention times previously established using unlabeled standards using Skyline software (MacCoss Lab, University of Washington, v23.1.0.455). Ratios of light to heavy peak areas were used for further analysis.

### Correlation analysis between peptides and metabolites

LiP-MS peptide residuals and abundances of identified metabolites were independently median-aggregated across replicates per condition and time point and mean-centered within each condition (glucose, nitrogen, and phosphate repletion). Analysis was restricted to peptides and metabolites significantly changing across time in at least one of the three repletion conditions. Pairwise Pearson correlations were computed between all significant LiP-MS peptides and metabolites, retaining only pairs with at least 8 paired non-missing observations. Resulting *p*-values were corrected for multiple testing using local false discovery rate estimation with fdrtool^91^ (v1.2.18). Significant correlations (R²) were clustered with Euclidean distances and Ward’s minimum variance linkage method. To explore functional context, peptides from proteins annotated to the GO Biological Process “translation” were analyzed as a subset.

### tRNA sequencing

#### Library construction

Samples before and after glucose (0, 1, 4, 10 min) or nitrogen repletion (0, 1, 2, 10 min) were prepared in duplicates. Frozen yeast pellets were resuspended in 650 µL of tRNA isolation buffer (100 mM NaOAc, pH 4.5, 10 mM EDTA, 1 % SDS) and mixed with 650 µL of 65°C hot acid phenol (pH 4.3). The cell suspension was vortexed vigorously and incubated at 65°C for 1 min, followed by addition of 100 µL BCP (Sigma Aldrich). Samples were centrifuged at 10,000 g and 4°C for 10 min and the aqueous phase was transferred to a new tube. After an additional round of hot acid phenol/BCP and a round of BCP-only extraction, RNA was precipitated from the aqueous phase by the addition of 2.5 volumes of ice-cold 100% ethanol and 25 µg glycogen (Ambion). After precipitation (for at least 30 min at –20°C), samples were centrifuged at 16,000 g and 4°C for 30 minutes, pellets were briefly air-dried and resuspended in RNase-free water. From these extracted RNA samples, tRNA sequencing libraries were generated following the previously published mim-tRNAseq method^46,92^, using Induro RT (New England Biolabs M0681L) for the reverse transcription instead of the commercially discontinued TGIRT. Libraries were quantified with the Qubit dsDNA HS kit and 120-bp single– end sequencing was performed on a NovaSeq 6000 platform.

#### Data analysis

Sequencing libraries were demultiplexed and adaptor-trimmed with Cutadapt^71^ with the following arguments: “--no-indels –q 30,30 –-trimmed-only –m 10”. Reads were further trimmed to remove low quality bases and the two 5′-RN nucleotides introduced in RT with “-j 40 –m 10 –u 2”. Given the low read depth of one replicate of nitrogen repletion at time 0, this sample was excluded in downstream analyses. Trimmed reads were aligned and analyzed with mim-tRNAseq^46^ (v1.3.11) with the arguments: “--species Scer –-cluster-id 0.90 –-deconv-cov-ratio 0.5 –-p-adj 0.01 –-max-mismatches 0.1 –-remap-mismatches 0.075 –-min-cov 0.0005 –remap –-misinc-thresh 0.1”.

### Polysome profiling by sucrose gradient

An aliquot of 300 µL of the same yeast cell lysate used for ribosome profiling was loaded onto 10-45% sucrose gradients in 20 mM Tris-HCl pH 8.0, 150 mM KCl, 5 mM MgCl_2_, and 0.5 mM DTT. Gradients were centrifuged at 38,000 rpm and 4°C for 1:40 hours using a WX 90 ultracentrifuge with a TH-641 rotor (Thermo Fisher Scientific). Absorbance at 260 nm along the sucrose gradient was measured.

### Fluorescent reporter assay

#### Plasmid and yeast strain construction

All reporter constructs were cloned using the integrative vector system EasyClone-MarkerFree^93^. Two analogous reporter constructs containing the monomeric fluorescent proteins yECitrine and mCherry were synthetically synthetized, with the same sequences used in Tunney et al.^29^, except the first threonine codon of yECitrine was changed from ACT to ACC to create an AgeI restriction cut site. Both proteins were designed under the ENO2 promoter followed by the full endogenous ENO2 5′ UTR for mCherry or the first four endogenous bases (AAGC) and a variable 5′ UTR sequence for yECitrine, as similarly described by May et al.^43^. These synthetic fragments were cloned into the AsiSI site of the EasyClone plasmids pCfB2903 (for mCherry, with HDR region integrating to the X chromosome, Addgene 73275) and pCfB2899 (for yECitrine, with HDR region integrating to the XI chromosome, Addgene 73271) by Gibson assembly, upstream of the ADH1 terminator. Additional variable 5′ UTR sequences were cloned into the yECitrine plasmid by digestion with AgeI and AvrII and Gibson assembly of designed synthetic sequences. As controls for low and high elongation time constructs, we also designed two constructs with a non-optimal (slow citrine, “citmax”) and optimal (fast citrine, “cit0”) citrine coding sequence, as designed in Tunney et al.^29^. All resulting plasmid sequences validated by Sanger sequencing are available in **Supplementary Data 1**.

Yeast strains were generated following instruction of the CRISPR-Cas9-based toolkit EasyClone-MarkerFree^93^. The YSBN6 strain was first transformed with the Cas9 expressing plasmid pCfB2312 (Addgene 78231) and selected in YPD plates with G418. Cas9-expressing cells were then transformed with the gRNA-expressing plasmid pCfB3020 (Addgene 73282) and the NotI-linearized mCherry construct and selected in YPD plates with G418 and nourseothricin. Integration was confirmed by detection of red fluorescence in the iBright system (Thermo Fisher Scientific) and colony PCR of the EasyClone X-2 locus. This parental strain was then grown in G418-only plates and replica plated in G418– and nourseothricin-containing plates to ensure removal of pCfB3020. This strain was then transformed with the gRNA-expressing plasmid pCfB3044 (Addgene 73286) and each of the NotI-linearized yECitrine constructs and selected in YPD plates with G418 and nourseothricin. Integration was confirmed by detection of yellow fluorescence in the iBright system and colony PCR of the EasyClone XI-2 locus. All generated strains are listed in **Supplementary Table 8**.

#### Flow cytometry

For each sample, 2 mL were collected in 2mL-microtubes, centrifuged (5,000 g for 5 min, same settings for all centrifugation steps), washed with 1 mL DPBS and centrifuged again. Cells were fixed with 1 mL 4% formaldehyde in DPBS for 15 minutes in the dark at room temperature, followed by two washes with DPBS. Samples were diluted 1:4 in DPBS and stored at 4°C until measurement. Flow cytometry was measured in an Attune NxT Cytometric Analyser (Thermo Fisher Scientific). Forward light-scatter (FSC) and side-scatter (SSC) were made with a 488-nm laser. Citrine fluorescence was measured with a 488-nm laser excitation and detected with a 495-nm long-pass optical filter followed by a 530/30 nm band-pass optical filter. mCherry fluorescence was measured with a 561-nm laser for excitation and a 600-nm long-pass optical filter followed by 620/15-nm band-pass optical filter. PMT values for each color channel were adjusted such that the mean of the non-fluorescent strain YSBN6 was 100. 50,000 events were collected for each sample.

Flow cytometry data was analyzed using the flowCore, flowViz, and flowStats R packages (see “Data and code availability”). A sequential two-step gating strategy was applied to isolate single, viable cells. First, debris was excluded using a rectangular gate on forward scatter area (FSC-A > 2·10^4^) and side scatter area (SSC-A > 1·10^4^). Second, doublets were removed by applying a “norm2Filter” on FSC height versus FSC area (scale factor = 3), retaining only singlets. Autofluorescence background was estimated as the median Citrine and mCherry in the wild-type strain (YSBN6) under glucose growth conditions and subtracted from all measurements. The Citrine-to-mCherry fluorescence ratio was calculated for each cell. For each construct, condition, and biological replicate, the median log_2_-transformed ratio was determined. Changes following glucose repletion were calculated by subtracting the median of the corresponding glucose-depleted sample within the same construct and biological replicate. Motif-associated effects were quantified as the difference between these baseline-normalized changes and those of the scrambled control. To estimate within-replicate sampling uncertainty, individual cells were resampled with replacement independently within each sample (1,000 bootstrap iterations), and the complete baseline-normalized contrasts were recalculated for each iteration. The 2.5th and 97.5th percentiles of the bootstrap distributions defined the 95% confidence intervals. The same bootstrap draws of the glucose-depleted samples were reused across contrasts involving the same baseline.

#### RT-qPCR

RNA was extracted using hot acid phenol (see details in “Total RNA sequencing”). Total RNA was treated with the DNA-free DNA Removal Kit (Invitrogen AM1906). For RT, 1 μg of DNase-treated RNA was reverse-transcribed using the Protoscript II first-strand cDNA synthesis kit (New England Biolabs E6560). RT-qPCR was performed with the KAPA SYBR Fast kit (Roche KK4611, **Supplementary Table 10**), with cDNA diluted 1:20 for Citrine and 1:5 for mCherry, and measured in technical triplicates in a Roche LightCycler 480 II (Roche Diagnostics). For analysis, ΔΔCt values for Citrine were calculated relative to mCherry. qPCR primer efficiency was tested with a serial dilution, with an efficiency of 1.9-2.1 for all primer pairs used.

#### Immunoblotting

Protein extraction was performed as previously described^94^. Cell pellets were resuspended in 200 µL protein extraction buffer (0.1 M NaOH, 0.05 M EDTA, 2% SDS, 1% β-mercaptoethanol), heated at 90°C with shaking (1,500 rpm) for 10 min, briefly centrifuged, acidified with 5 µL 4 M acetic acid, vortexed for 30 s, and reheated at 90°C with shaking for an additional 10 min. After centrifugation, 50 µL loading buffer (0.25 M Tris-HCl pH 6.8, 50% glycerol, 0.05% bromophenol blue) was added. Samples (15 µL) were loaded on NuPAGE 4–12% Bis-Tris gels (1.0 mm, 15-well, Invitrogen NP0323BOX) using Bolt MES running buffer (Invitrogen B0002). Duplicate gels were run in parallel for separate detection of citrine and mCherry.

Proteins were transferred to nitrocellulose (Amersham 10600015) by semi-dry transfer using freshly prepared 1x transfer buffer (48 mM Tris, 39 mM glycine, 0.75% SDS, 10% ethanol) at 20 V for 25 min. Membranes were rinsed with water, stained with Ponceau S (0.5% Ponceau S, 1% acetic acid) for 10 min, rinsed with water and imaged. After destaining with PBST (PBS with 0.1% Tween-20), membranes were blocked in 5% non-fat dry milk in PBST for 1 h at room temperature. Primary antibodies were diluted in blocking buffer (anti-GFP at 1:5000, Origene TP401, and anti-mCherry at 1:2000, Novus Biologicals NBP1-96752) and incubated overnight at 4°C with gentle agitation. Membranes were rinsed and washed twice for 10 min each in PBST. HRP-conjugated secondary antibodies (anti-Rabbit, Dianova 111-035-003, and anti-Mouse, Dianova 115-035-003) were diluted in blocking buffer and incubated for 60 min at room temperature, followed by three 10-min washes in PBST and a final rinse in PBS. Membranes were incubated with ECL substrate (SuperSignal Pico, Thermo Fisher Scientific 34577) for 5 min protected from light, and imaged using the iBright system (Thermo Fisher Scientific).

### Quantification and statistical analysis

All details of the statistical analyses (*p*-value, value of *n*, contrast analyses) can be found in the “Results” section and the figure legends.

## SUPPLEMENTARY INFORMATION

**Supplementary Table 1. Proteomic and metabolic characterisation of nutrient depletion and repletion.** (A) Differential protein abundance of glucose-, nitrogen– and phosphate-depleted cultures compared to cells grown in SD medium (*limma*). Columns: log2 fold change (logFC), p-value and Benjamini-Hochberg adjusted p-value for each depletion (G, N, P), and gene name. (B) GO Biological Process enrichment (clusterProfiler) of significantly up– and down-regulated proteins in each nutrient depletion. (C) Normalised (internal standards and probabilistic quotient normalisation) and batch-corrected abundances of all untargeted metabolic features (rows, ‘m/z retention time’) in all samples (columns, condition + time point + replicate). (D) Differential abundance of metabolic features along the 30-min glucose repletion (*limma*, cubic spline with knots at 0.5, 2.5 and 5 min). Columns: spline coefficients (X1-X4), average expression, F statistic, p-value, adjusted p-value, top compound annotation (name, class, adduct, score, source) and mean abundance per time point. (E) Same as D, for the nitrogen repletion. (F) Same as D, for the phosphate repletion. (G) Peak set enrichment analysis (mummichog, MetaboAnalystR) of metabolic pathways for each time-pattern cluster of significantly changing features (Fisher’s exact test p < 0.05). Columns: nutrient, cluster, KEGG pathway, Fisher’s exact test p-value (FET) and enrichment factor.

**Supplementary Table 2. Spike-in-calibrated ribosome profiling and total RNA sequencing upon nutrient repletion.** (A) Differential mRNA abundance along the glucose repletion (*limma-voom*, cubic spline with knots at 0.5, 2.5 and 5 min). Columns: spline coefficients (X1-X4), average expression, F statistic, p-value, adjusted p-value, TPM per time point and gene name. (B) Same as A, for the nitrogen repletion. (C) Same as A, for the phosphate repletion. (D) Differential ribosome footprints along the glucose repletion (*limma-voom* with RUVSeq normalisation, cubic spline). Columns as in A. (E) Same as D, for the nitrogen repletion. (F) Same as D, for the phosphate repletion. (G) Absolute ribosomes-per-transcript per gene and sample, calibrated with the Fluc (trisome) and Rluc (disome) spike-ins. (H) Differential ribosomes-per-transcript along the glucose repletion (*limma* on log-transformed ribosomes-per-transcript, cubic spline). Columns: spline coefficients (X1-X4), average expression, F statistic, p-value, adjusted p-value, ribosomes-per-transcript per time point and gene name. (I) Same as H, for the nitrogen repletion. (J) Same as H, for the phosphate repletion. (K) GO Biological Process enrichment (clusterProfiler) of each time-pattern cluster of transcripts with significantly changing ribosomes per transcript. (L) A-site occupancies of all codons for short (20-23 nt) and long (27-32 nt) footprints per sample. (M) Elongation-time estimates (log_2_ geometric mean of the A-site occupancies of all codons of each gene) for short (20-23 nt) and long (27-32 nt) footprints per sample, and initiation estimate (z-scored mean YFP of the 5’UTR MPRA of Akirtava et al.) per gene.

**Supplementary Table 3. Performance of RiboNet models of ribosome footprint counts and distribution.** (A) Benchmark of the full RiboNet model, its ablated versions (5’UTR ablation, 3’UTR ablation, CDS only, counts-only training) and linear baselines (RNA abundance, codon usage) over 50 training iterations with different train/validation/test splits. (B) Spearman correlation between predicted and observed per-codon footprint distribution of test-set genes, per repletion, time point and footprint length, and Spearman correlation between biological replicates.

**Supplementary Table 4. tRNA abundance, aminoacylation and condition-specific codon decoding**. (A) Abundance of cytoplasmic tRNA anticodon families (% of cytoplasmic tRNA reads) per sample, for the glucose and nitrogen repletions. (B) Aminoacylation (% of 3’-CCA-ending reads) of cytoplasmic tRNA anticodon families per sample. (C) Targeted metabolomics of amino acids: light-to-heavy ratios against ^13^C^15^N-labelled internal standards per sample.

**Supplementary Table 5. Protein abundance and structural changes of the proteome upon nutrient repletion by LiP-MS.** (A) Differential protein abundance along the glucose repletion (*limma*, cubic spline with knots at 0.5, 2.5 and 5 min). Columns: spline coefficients (X1-X4), average expression, F statistic, p-value, adjusted p-value, gene name and mean abundance per time point. (B) Same as A, for the nitrogen repletion. (C) Same as A, for the phosphate repletion. (D) Differential structural changes (LiP-MS residuals) along the glucose repletion (*limma*, cubic spline). Columns: spline coefficients, average expression, F statistic, p-value, adjusted p-value, peptide annotation, and mean residual per time point. (E) Same as D, for the nitrogen repletion. (F) Same as D, for the phosphate repletion. (G) Significant Pearson correlations (local FDR < 0.05, fdrtool) between significantly changing LiP-MS peptides and significantly changing identified metabolites across all 24 matched condition-time points. Columns: peptide ID, metabolite ID, p-value, q-value, local FDR, Pearson R (column named R2), peptide annotation (gene_start-end), metabolite annotation and GO ‘translation’ membership. 9,352 of 1,429,866 tested pairs; the full table is available on Zenodo.

**Supplementary Table 6. Identification and characterization of the 5’UTR GAUY motif.** (A) GAUY motifs identified from the attribution scores of footprint counts (TF-MoDISco). Columns: gene, motif attribution score and location per condition, detected conditions, gene average ribosomes-per-transcript, distance from start codon, distance from longest TSS (TSS) and consensus TSS (cons), presence of upstream AUGs. (B) Evolutionary conservation of high-confidence GAUY seqlets: Simple Enrichment Analysis (SEA, MEME suite) of each gene’s seqlet among the 200-nt upstream sequences of its fungal homologs at increasing evolutionary distances. Columns: number of homologs, enrichment ratio of motif, p-value, median enrichment ratio of random k-mers, corrected enrichment (motif vs k-mers), evolutionary distance. (C) Over-representation analysis of GO terms (Biological Process, Cellular Component, Molecular Function) among high-confidence GAUY-containing genes, using all modelled genes as universe. (D) Comparison of codon frequency between high-confidence GAUY-containing genes vs non-GAUY-containing genes for all 61 codons. Statistical differences are determined by U Mann-Whitney test and FDR-corrected using Benjamini-Hochberg.

**Supplementary Table 7. Media composition for glucose, nitrogen and phosphate repletion.**

**Supplementary Table 8. YSBN6-derivative strains for reporter assay.**

**Supplementary Table 9. Comparison of features deployed by deep learning models of mRNA translation.** Abbreviations: RPF, ribosome footprint; CNN, convolutional neural network; FNN, feed-forward neural network; RNN, recurrent neural network.

**Supplementary Table 10. Primers for library construction and RT-qPCR.**

**Supplementary Data 1. Sequences and annotation of all plasmid sequences used for strain transformation (GenBank file).**

